# Regulation of large-conductance Ca^2+^- and voltage-gated K^+^ channels by electrostatic interactions with auxiliary β subunits

**DOI:** 10.1101/2021.02.22.432338

**Authors:** Yutao Tian, Stefan H. Heinemann, Toshinori Hoshi

## Abstract

Large-conductance Ca^2+^- and voltage-gated K^+^ (BK K_Ca_1.1) channel complexes include pore-forming Slo1 α subunits and often auxiliary β subunits, latter of which noticeably modify the channel’s pharmacological and gating characteristics. In the absence of intracellular Ca^2+^, β1 and β4 modestly shift the overall voltage dependence of the channel to the positive direction by decreasing the probability that the ion conduction gate is open without any allosteric influence from the channel’s voltage or Ca^2+^ sensors. This intrinsic open probability is also critically regulated by the intracellular-facing ^329^RKK^331^ segment of human Slo1 (hSlo1) downstream of the transmembrane segment S6 in association with two negatively charged residues in S6 (E321 and E324) (Tian et al., Proc Natl Acad Sci USA, 116, 8591-8596, 2019). This study examined how β1/β4 and the RKK segment function together to control the channel gate. With select mutations in the RKK segment, inclusions of β1 or β4 can dramatically increase the intrinsic gate opening probability and shift the overall voltage dependence of the channel to the negative direction by up to 200 mV without Ca^2+^. This remarkable shift is mediated at least in part by electrostatic interactions between the Slo1 RKK and β N-terminal segments as suggested by the results of double-mutant cycle analysis, ionic strength experiments, and molecular modelling. With or without auxiliary β subunits, the Slo1 RKK and E321/E324 segments are thus critical determinants of the intrinsic open probability of the ion conduction gate and changes in the electrostatic environment near the RKK-EE segments are a potential mechanism of pharmacological gating modifiers.

## Introduction

Large-conductance Ca^2+^- and voltage-gated K^+^ (BK) channel complexes allow K^+^ flux in response to membrane depolarization and/or an increase in intracellular Ca^2+^ concentration ([Ca^2+^]_i_), generally limiting cellular electrical excitation (1). To accomplish this task, activation of the transmembrane voltage sensors and intracellular divalent cation sensors of the tetrameric pore-forming Slo1 protein is allosterically coupled to opening of the ion conduction gate (Fig. 1A) (2, 3). The functional interactions among the gate and the sensors are well described by the Horrigan and Aldrich (HA) model of Slo1 gating (4).

**Figure 1.**
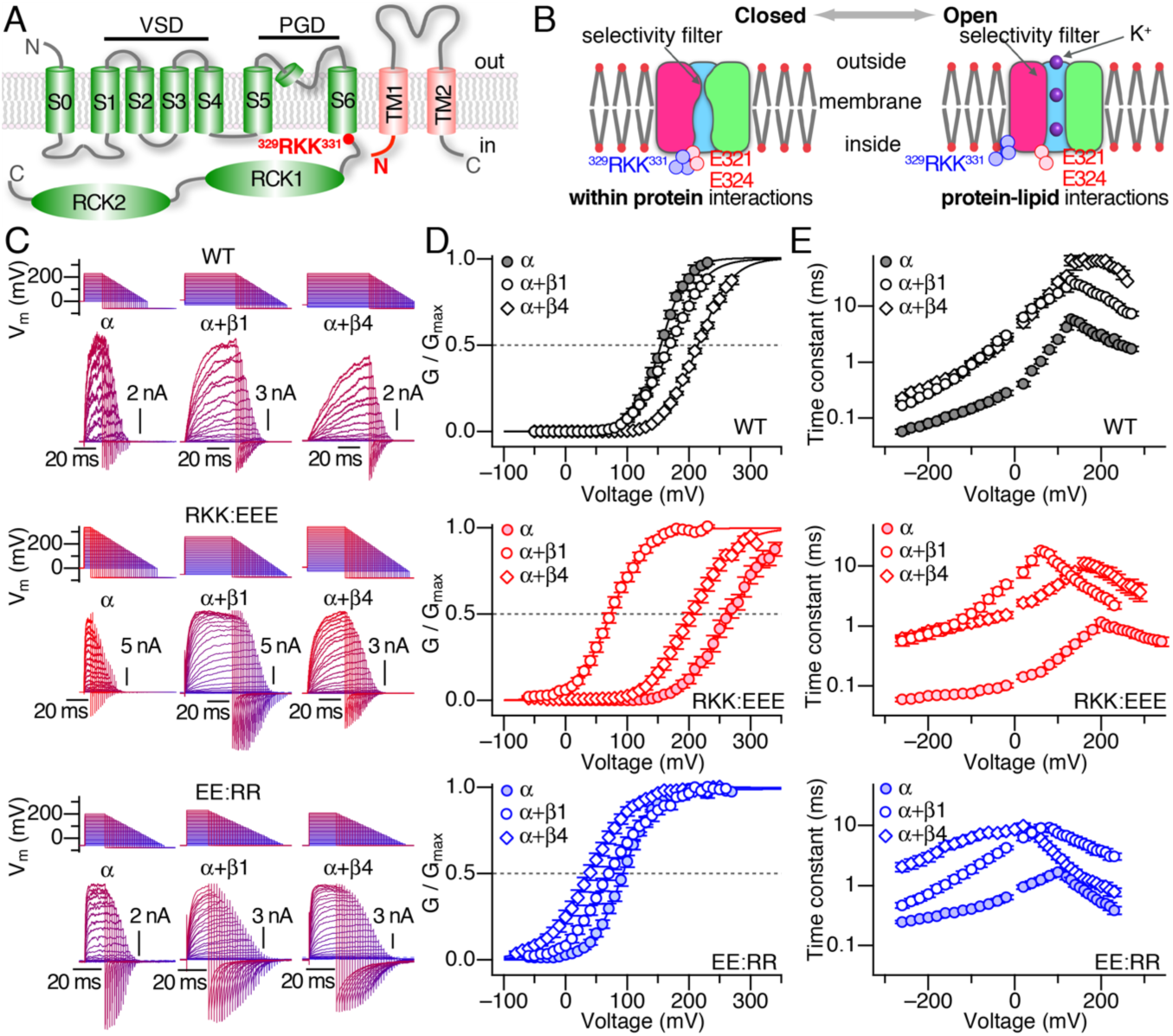
Contrasting effects of coassembly with β1/4 in wild-type hSlo1 and hSlo1 RKK-EE ring mutants. **A**. Schematic organizations of one pore-forming hSlo1 (*left*, green) subunit and a β subunit (*right*, pink). One Ca^2+^ sensor site exists in each of the two RCK domains in one subunit. Not drawn to scale. PGD: pore-gate domain. VSD: voltage-sensor domain. The two areas important for this study are shown in red (hSlo1 ^329^RKK^331^, hSlo1 E321, hSlo1 E324, and *β* N terminus). **B**. Schematic illustrations of the interactions made by ^329^RKK^331^ and E321/E324 in the closed state (*left*) and the open state (*right*). Only one of the four RKK-EE interactions are shown. Small red circles represent lipid oxygen atoms. Not drawn to scale. The cytoplasmic area is not shown. **C**. Illustrative current traces from α alone (*left*), with β1 (*center*), and with β4 (*right*). The voltage and current sweeps use the same color scheme, from hyperpolarized (blue) to depolarized (red) test voltages. **D**. Normalized GV curves from α alone (filled circles), with β1 (open circles), and with β4 (open diamonds). Smooth curves are Boltzmann-type fits (see Tables S1 and S2). **E**. Time constants of ionic current relaxation at different voltages from *α* alone (filled circles), with β1 (open circles), and with β4 (open diamonds). In each panel in **C**-**E**, the results from three pore-forming hSlo1 constructs are shown. *Top*: wild-type (WT) hSlo1, *middle*: hSlo1 R329E:R330E:R331E (RKK:EEE), *bottom*: hSlo1 E321R:E324R (EE:RR). All results were obtained without Ca^2+^.

Slo1 BK channels exist nearly ubiquitously in different tissue and cell types, and contribute to multitudes of physiological processes (1). This functional versatility in vertebrates is conferred often by coassembly with auxiliary β and γ (LRRC) subunits in a tissue-dependent manner (5, 6). For example, Slo1 complexes containing β1 subunits (Slo1+β1; α+β1) noticeably contribute to regulation of smooth muscle tone, and Slo1+β4 (α+β4) channels participate in neuronal function (6). Coassembly with β subunits alters the pharmacology of the channel as well as multiple aspects of gating (1, 6). For example, dehydrosoyasaponin-1, a plant triterpene glycoside, more effectively opens Slo1+β1 than Slo1 alone (7). An apparent increase in the overall sensitivity to intracellular Ca^2+^ in Slo1+β1 is also well documented (5, 6). In the absence of intracellular Ca^2+^, inclusion of four β1 subunits shifts the overall voltage dependence of activation slightly to the positive direction and inclusion of β4 subunits produces a more noticeable shift, >20 mV (1, 6). Further, activation and deactivation kinetics of Slo1+β1 and Slo1+β4 without Ca^2+^ are markedly slower than those of Slo1 alone (1, 6). The aforementioned gating changes and others induced by β1 and β4 have been characterized using the HA model (8-10). While many parameters of the gating model change (9-11), one principal effect of coassembly with β1 or β4 is to decrease the probability that the ion conduction gate is open without activation of the voltage or divalent cation sensors (9, 10) as specified by the equilibrium constant L_0_ in the HA model (4). The structural basis for the decrease in L_0_ is unclear. In contrast with β subunits, inclusion of γ1 (LRRC26) strengthens coupling between the ion conduction gate and the voltage sensors (D in the HA model) and drastically moves the overall voltage dependence of activation without Ca^2+^ to the negative direction by up to ∼135 mV (12).

Each β subunit possesses two short transmembrane domains, TM1 and TM2, connected by a variable extracellular loop; both N- and C-terminal ends face the intracellular side (Fig. 1A) (1, 6, 13). Atomic-level structures of the β4-containing human Slo1 (hSlo1) channel are available (13). Largely consistent with the results of an earlier cross-linking study (14), each of the four β4 subunits is located between two neighboring voltage-sensor domains (VSDs) of the tetrameric pore-forming hSlo1 or α complex, just peripheral to the central pore-gate domain (PGD) comprised of S5, P, and S6 segments (Fig. 1A), where membrane lipids would otherwise occupy (13). It is very probable that hSlo1+β1 are organized in a similar way (13). It is thought that the presence of β subunits does not induce a new conformation in the pore-forming subunits but stabilizes select pre-existing conformations found without β (13).

Some amino-acid residues within β1 important for their effects on channel gating are known. While multiple structural components of β1 and β4 subunits may contribute to the slower kinetics in β1- and β4-containing hSlo1 channels, the N terminus may be particularly important (13). Two distal N-terminal Lys residues, K4 and K5, functionally interact with the Slo1 voltage sensors (15). R11 and C18 in β1 potentiate the consequence of the interaction between omega-3 docosahexaenoic acid (DHA) and Y318 (using the human Slo1 (hSlo1) numbering in AAB65837) in S6 of the PGD to increase L_0_ (16, 17). Activation of the intracellular Ca^2+^ sensors may be also coupled to E13 and T14 in the β1 N terminus to change the apparent voltage dependence with Ca^2+^ present (18). Functional coupling between β1 and pore-forming Slo1 subunits also depends on *S*-acylation of select Cys residues within Slo1 (19).

We recently showed that the sequence ^329^RKK^331^ in hSlo1 immediately downstream of S6 in the primary structure (Fig. 1A) is a critical determinant of the channel gating, especially of the intrinsic behavior of the ion conduction gate as specified by L_0_ in the HA model (20). The closed conformation of the gate is stabilized by ion-ion interactions between the RKK segment in one subunit and S6 Glu residues (E321 and E324) in a neighboring subunit of the channel (Fig. 1B), forming a ring-like inter-subunit interaction network termed the RKK-EE ring (20). In contrast, the open conformation is stabilized in part by ion-dipole interactions between the RKK segments and membrane phospholipid oxygen atoms (Fig. 1B) (20); the lipid molecules surrounding the hSlo1 channel may have a role of as an integral component of the channel’s gating machinery (13, 20, 21).

Here we investigated the interactions between the RKK-EE ring and auxiliary β1/β4 subunits, both of which are known to alter the behavior of the hSlo1 ion conduction gate characterized by the equilibrium constant L_0_. In the hSlo1+β4 structure, the RKK segment is found near the N terminus of β4 (13). Previous studies have established that without intracellular Ca^2+^, coassembly of hSlo1 with β1 or β4 shifts the half-activation voltage (V_0.5_) to the positive direction (9, 10). We show in contrast that, when the ion-ion interactions forming the RKK-EE ring is impaired, coassembly with β1 or β4 moves the voltage dependence of the resulting channels to the negative direction, by up to 200 mV by increasing L_0_. This β1/β4-dependent increase in L_0_ involves an electrostatic interaction between the N terminus of β1/β4 and the RKK segment, preventing the formation of the closed-state stabilizing RKK-EE ring. The results reinforce the fundamental importance of the RKK-EE ring, capable of interacting with membrane lipids and also with β subunits, in Slo1 channel gating, and suggest that the electrostatic environment around the RKK-EE ring may be a potential pharmacological target of gating modifiers.

## Materials and methods

### Channel expression

Ion channel genes were expressed in human embryonic kidney (HEK) tsA cells obtained from two sources (Prof. R. Horn, Jefferson University, Philadelphia, PA and Sigma-Aldrich #96121229) as described previously (20). Plasmid DNAs coding for hSlo1 (AAB65837 in pCI-neo) and human β1 (AAH25707) or β4 (KJ893642) were transfected into the cells with either FuGene6 (Promega) or GenJet v. II (SignaGen) according to the vendor instructions. The cells were used for electrophysiological measurements ∼36 to 48 h after transfection. The β constructs used have green fluorescent protein (GFP) fused to the C terminus (16). The results similar to those in Fig. 1 were also obtained using β1 and β4 without the fused GFP tag. For coexpression of hSlo1 and β subunit genes, the hSlo1 and β plasmid DNA weight ratio was 1:2. For coexpression of hSlo1 and γ1 subunit genes, the DNA weight ratio was 1:1. In the experiments with γ1, GFP (pcDNA3) was used as the transfection marker. Functional coassembly with β1/4 subunits in each patch was inferred by the characteristic slowing of the activation/deactivation kinetics induced by β subunits (16) (Figs. S12-S14). In hSlo1+γ1, functional inclusion of γ1 was inferred by the left-shifted voltage dependence of activation (12). Mutations in hSlo1 and β1/4 were introduced by PCR-mediated mutagenesis (NEB) using oligonucleotide primers (IDT) and verified by sequencing (DNA Sequencing Facility, University of Pennsylvania Genetics).

### Cell culture

Cells were grown in Dulbecco’s Minimal Eagles Medium (DMEM) with glutamine and 4.5 g/L glucose (Mediatech) supplemented with fetal bovine serum (10%; Sigma-Aldrich) and penicillin-streptomycin (0.1 U/L; ThermoFisher) in a humidified 37 °C incubator with 5% CO_2_.

### Electrophysiology

The electrophysiological data collection and analysis protocols using the inside-out configuration of the patch-clamp method at room temperature were as described recently in detail (20). The standard “extracellular” solution contained (in mM):140 KCl, 2 MgCl_2_, and 10 HEPES, pH 7.2 with *N*-methyl-*D*-glucamine (NMG). The Ca^2+^-free/no added Ca^2+^ “intracellular” solution contained (in mM): 140 KCl, 11 EGTA, and 10 HEPES, pH 7.2 with NMG. The intracellular solution with 100 µM Ca^2+^ did not contain any Ca^2+^ chelator. All salts were obtained from Sigma-Aldrich. The ionic strength experiments were performed using an agar-bridge (15%, 2000 mM KCl) ground electrode. The results were analyzed with IgorPro (v. 6 and 8, WaveMetrics).

### HA model-based fitting

Ionic current properties in the absence of Ca^2+^ were simulated using the HA model (4, 22) in the following manner. For hSlo1, the parameter values were taken from our previous study (20). For hSlo1+β1, the parameter values for hSlo1 were first altered according to the adjustments made by Orio and Latorre to explain the gating differences between hSlo1 and hSlo1+β1 (9). The value of L_0_ was then fine-tuned based on the single-channel results at negative voltages without Ca^2+^ as presented here, and the value of the coupling factor D was adjusted by eye so that the simulated GV curve matches the experimentally observed curve (Fig. S1). For hSlo1+β4, the parameters were adjusted as for hSlo1+*β*1 so that the simulated and measured GV curves match (Fig. S1). For hSlo1 RKK:EEE+β1 and hSlo1 RKK:EEE+*β*4, the parameters for hSlo1 RKK:EEE (20) were adjusted as described above for those with wild-type hSlo1 (Fig. S5).

### Double mutant cycle analysis

The coupling factor (23) is defined here as Ω = [ L_0 (hSlo1+β1)_ * L_0 (hSlo1 RKK + β1 AQKR:XXXX)_] / [ L_0 (hSlo1 RKK:EEE+β1)_ * L_0 (hSlo1+ β1 AQKR:XXXX)_], where X is the amino acid substituted (R, G, V, T, A, Q, or E). For example, L_0 (hSlo1+ β1)_ represents the equilibrium constant L_0_ according to the HA model for hSlo1+β1. In the experiments examining the mutations hSlo1 RKK:EEE and β1 AQKR:EEEE, the L_0_ values were directly estimated from single-channel openings without Ca^2+^ at negative voltages, at least 120 mV more negative than the respective macroscopic V_0.5_ values. The L_0_ voltage dependence (z_L_ in the HA model) was estimated from the partial charge movements associated with the voltage dependence of activation and deactivation kinetics at extreme positive and negative voltages (20). For other β1 mutants, the L_0_ values were estimated by fitting macroscopic GV curves in the following way. Starting from the published parameter values (20), the z_L_ values were determined from the voltage dependence of activation and deactivation, and only the L_0_ value was manually adjusted so that the simulated GV curves match the experimental results as judged by eye (Fig. S9).

### Structural models of hSlo1

Molecular dynamics (MD)-based energy minimizations were utilized to infer the structural features of the following channels embedded in membranes, without divalent cations and with both Ca^2+^ and Mg^2+^ fully bound to the respective ion sensors: hSlo1, hSlo1 RKK:EEE, hSlo1+β1, hSlo1+β4, hSlo1 RKK:EEE+β1, and hSlo1 RKK:EEE+β4. The starting models of hSlo1 were from PDB# 6V3G (divalent cation free) and PDB# 6V38 (Ca^2+^ and Mg^2+^ bound), and those of hSlo1+β4 were from PDB# 6V35 (divalent cation free) and PDB# 6V22 (Ca^2+^ and Mg^2+^ bound) (https://opm.phar.umich.edu). Missing and unresolved residues were filled with SWISS-MODEL (https://swissmodel.expasy.org) using the default options (24). The structures of hSlo1+β1 was modelled from those of hSlo1+β4 with SWISS-MODEL (https://swissmodel.expasy.org) (24). The resulting models contained residues 22 through 1056 of hSlo1, residues 9 through 191 of β1, and residues 8 through 205 of β4. Water (TIP3P; S1 and S3) and K^+^ ions (S0, S2, and S4) were manually placed in the selectivity filter. Each structure was embedded in a bilayer membrane with 150 mM or 2000 mM KCl using CHARMM-GUI (25, 26) as described (20). To better correlate with the electrophysiological results, the membrane lipid composition was designed to roughly mimic a native cell membrane (27). The outer leaflet contained: cholesterol (20%), 1,2-dipalmitoyl-sn-glycero-3-phosphocholine (DPPC, 40%), and 1,2-dioleoyl-sn-glycero-3-phosphocholine (DOPC, 40%). The inner leaflet contained: cholesterol (20%), DPPC (25%), DOPC (25%), 1,2-dipalmitoyl-sn-glycero-3-phosphoethanolamine (DPPE, 10%), 1,2-dioleoyl-sn-glycero-3-phosphoethanolamine (DPPE, 10%), 1,2-dipalmitoyl-sn-glycero-3-phosphoserine (DPPS, 5%), and 1,2-dioleoyl-sn-glycero-3-phospho-L-serine (DPPS, 5%). We showed in our previous study (20) that the lipid composition does not influence the interactions between the RKK and E321/E324 segments. A typical system contained 600,000 atoms all together. The equilibration and simulation runs were performed with CUDA-enabled NAMD2 (28) (http://www.ks.uiuc.edu/) running on Mac OS or Ubuntu OS using the CHARMM36m force field under the periodic boundary NPT condition at 37 °C with the default simulation parameter values suggested by CHARMM-GUI. Each simulation run was ≥120 ns and the structures were examined with VMD (v1.9; http://www.ks.uiuc.edu/) and PyMol (v2.4; Schrödinger), and visualized by PyMol. The root-mean-square deviation (RMSD) values were stable within ∼20 to 50 ns. The structural images shown are from the last ns of the simulations.

### Data descriptors

Unless otherwise noted, statistical results are shown as means ± standard deviation (n), where n represents the number of independent measurements. Fit parameters are shown as means ± 95% confidence interval as calculated in IgorPro.

## Results

### Functional consequences of hSlo1 coassembly with *β*1 or *β*4

In the virtual absence of Ca^2+^, currents through wild-type hSlo1 channels fully activated within a few ms at the positive voltages where the macroscopic conductance is saturated (>200 mV; Fig. 1C *top left*; Fig. 1D, E *top* filled black circles). On repolarization to the negative voltages where the conductance is very small (e.g., –200 mV), the currents decayed rapidly with a time constant of < 0.1 ms (Fig. 1E *top* filled black circles). At steady state, the half-activation voltage (V_0.5_) of the wild-type hSlo1 channel was about 160 mV (Fig. 1D *top* filled black circles).

Confirming numerous previous results (5, 6), coassembly with β1 slowed ionic current kinetics at all voltages ∼8 fold (Fig. 1C *top middle*; Fig. 1E *top* open circles). In addition, the normalized macroscopic conductance-voltage (GV) curve was shallower and its V_0.5_ slightly moved to the positive direction compared with the mean V_0.5_ value in hSlo1 (Fig. 1D *top* open circles; ΔV_0.5_ = 13.4 ± 5.5 mV (35)). In hSlo1+*β*4, the slowing of the activation kinetics at positive voltages was much more noticeable (Fig. 1C *top right*; Fig. 1E *top* open diamonds) and the shift of V_0.5_ to the positive direction was markedly greater (Fig. 1D *top* open diamonds, ΔV_0.5_ = 61.4 ± 8.1 mV (36)).

The aforementioned changes in V_0.5_ and current relaxation kinetics by coassembly with *β*1 and *β*4 involve alterations of multiple parameters in the HA model of Slo1 gating; however, one main consequence is a near 10-fold decrease in L_0_ (9, 10). The decreases in L_0_ largely account for the positive shifts of V_0.5_ by β1 and β4; without such decreases in L_0_, the V_0.5_ values would shift to the negative direction (Fig. S1). We demonstrated earlier that the same parameter L_0_ is also controlled by ^329^RKK^331^ located in the cytoplasmic domain just C terminal to S6 (Fig. 1A) (20). When the ion conduction gate is closed, each RKK segment interacts with E321 and/or E324 within S6 in a neighboring subunit (Fig. 1B). With the gate open, the RKK segments interact with nearby membrane lipid oxygen atoms (Fig. 1B). The hSlo1 triple charge-reversal mutation R329E:K330E:K331E (RKK:EEE) disrupts both the RKK-EE ring stabilizing the closed conformation and the RKK-lipid oxygen interactions stabilizing the open conformation. The net effect is that the equilibrium constant L_0_ between the closed and open conformations of the ion conduction gate increases slightly by about 3-fold (20). Coupled with a probable change in the voltage sensor equilibrium (20), the overall voltage dependence shifts to the positive direction (Fig. 1D, E *middle* filled red circles compared with the filled black circles in Fig. 1D *top*). The Slo1 mutation E321R:E324R (EE:RR) impairs the closed-state-stabilizing interactions and shifts V_0.5_ to the negative direction (Fig. 1C, D-E *bottom* filled blue circles).

### hSlo1 mutants with impaired RKK-EE rings coassembled with *β*1 or *β*4

Both the hSlo1 RKK-EE ring (20) and auxiliary β subunits (9, 10) regulate the equilibrium constant L_0_. Thus, we examined how coassembly with β1 or β4 alters the electrophysiological properties of the channels containing mutations in the RKK-EE rings. Coexpression of hSlo1 RKK:EEE with β1 or β4 markedly slowed the current relaxation kinetics at all voltages (Fig. 1E *middle* red open circles and diamonds) essentially as observed for wild-type hSlo1 (Fig. 1E *top*, black open circles and diamonds), signifying that the mutant hSlo1 channel retained the ability to coassemble with β1 or β4. However, the changes in V_0.5_ caused by coassembly of hSlo1 RKK:EEE with β1 or β4 were clearly different from those observed with wild-type hSlo1. Unlike with wild-type hSlo1 where β1 only modestly moved the V_0.5_ to the positive direction (Fig. 1D *top* black open circles), coassembly of hSlo1 RKK:EEE with β1 moved the V_0.5_ to the negative direction to 74 mV or ΔV_0.5_ =∼ –192 mV (Fig. 1C, D *middle* red open circles). Coassembly with *β*4 also moved the V_0.5_ to the negative direction (V_0.5_ = 205 mV, ΔV_0.5_ = –60 mV; Fig. 1C, D *middle*). However, the shift by β4 was clearly smaller than that by β1, in contrast with the results with wild-type hSlo1 where the effect of β4 on V_0.5_ was much greater than that of *β*1 (Fig. 1D *middle* vs. *top*).

### Other hSlo1 RKK mutations with β1 or β4

The RKK segment of hSlo1 was mutated to other amino acids and the functional properties of the resulting mutant channels were examined with β1 or β4. The GV and kinetic properties of hSlo1 RKK:DDD+β1 and hSlo1 RKK:DDD+β4 (Fig. S2 *first row*) closely resembled those of hSlo1 RKK:EEE+β1 and hSlo1 RKK:EEE+β4 (Fig. 1C-D *middle*). For instance, coassembly of hSlo1 RKK:DDD with β1 produced a much greater negative shift of V_0.5_ than *β*4, –198 vs. –78 mV, respectively (Fig. S2A; Fig. S2B). The negative V_0.5_ shifts by β1 and β4 were considerably smaller with the neutral but polar mutation hSlo1 RKK:QQQ (Fig. S2 *second row*). With hSlo1 RKK:AAA, the negative V_0.5_ shift by β1 was even smaller and β4 produced a small positive V_0.5_ shift (Fig. S2 *third row*). With hSlo1 RKK:VVV, β1 had almost no effect and β4 produced a noticeable positive shift of V_0.5_ (Fig. S2 *bottom row*). These mutagenesis results, changing RKK to different amino acids, together suggest that the electrostatic characteristics of the hSlo1 RKK segment may be an important determinant in how β1 and β4 regulate the channel’s V_0.5_. This possibility is better suggested when the V_0.5_ values of the mutant hSlo1+β1 and hSlo1+β4 complexes are plotted as a function of the hydrophobicity index (29) of the substituted amino acid (Fig. S3). It is unclear why the results from the wild-type channel with RKK do not appear to fit the overall trend.

The mutation E321R:E324R (EE:RR) also impairs the RKK-EE ring and destabilizes the closed conformation of the gate (Fig. 1B), moving the V_0.5_ to the negative direction (Fig. 1C *bottom*) (20). Coexpression of hSlo1 EE:RR with β1 or β4 shifted the V_0.5_ to the negative direction (Fig. 1D *bottom*) albeit not as much as that found with hSlo1 RKK:EEE. The impacts of coassembly of hSlo1 EE:RR with β1 and β4 on the ionic current time course were also markedly different from those seen with wild-type hSlo1. With wild-type hSlo1, inclusion of β1 or β4 slowed the kinetics at all voltages (Fig. 1E *top*). In contrast, with EE:RR, β1 preferentially slowed the activation time course at positive voltages, consistent with preferential stabilization of the closed conformation of the gate. β4 selectively slowed down the deactivation kinetics at negative voltages, indicating preferential open state stabilization (Fig. 1E *bottom*). Qualitatively similar results were obtained with hSlo1 EE:QQ and hSlo1 EE:AA (Fig. S4).

### Coassembly with *β* subunits increases L_0_ in RKK:EEE

Coassembly of *β*1/*β*4 with wild-type hSlo1 alters the ion conduction gate behavior in the absence of any allosteric influence (9, 10), which could be experimentally estimated from single-channel measurements at extreme negative voltages without Ca^2+^. Illustrative single-channel openings from membrane patches containing multiple hSlo1 RKK:EEE, hSlo1 RKK:EEE+β1, and hSlo1 RKK:EEE+β4 channels at –200 mV, more than –250 mV away from the respective GV V_0.5_ values, are depicted in Fig. 2A. Under such a condition, the P_o_ value for hSlo1 RKK:EEE was 2.9 x 10^-5^ (Fig. 2C filled circles), which is ∼10-fold greater than that for wild-type hSlo1 at –120 mV (about –280 mV away from its V_0.5_) (Fig. 2B filled circles). With wild-type hSlo1, inclusion of β1 decreased P_o_ by about ∼10-fold (Fig. 2B, D). In contrast, with hSlo1 RKK:EEE, β1 increased P_o_ at –200 mV by ∼40-fold and β4 increased P_o_ by ∼10-fold (Fig. 2C, D). The conspicuous increases in P_o_ at the extreme negative voltages without Ca^2+^ correspond to similar increases in L_0_ in the HA model, about –9 and –6 kJ/mol, and these L_0_ changes primarily determine the shifts in V_0.5_ by *β*1 and β4 in hSlo1 RKK:EEE according to the HA model framework (Fig. S5). Without the large increases in L_0_, the V_0.5_ changes are difficult to explain (Fig. S5). Thus, the changes in V_0.5_ observed here by coassembly with β subunits mostly reflect alterations in one gating step, that between the closed and open conformations of the ion conduction gate.

**Figure 2.**
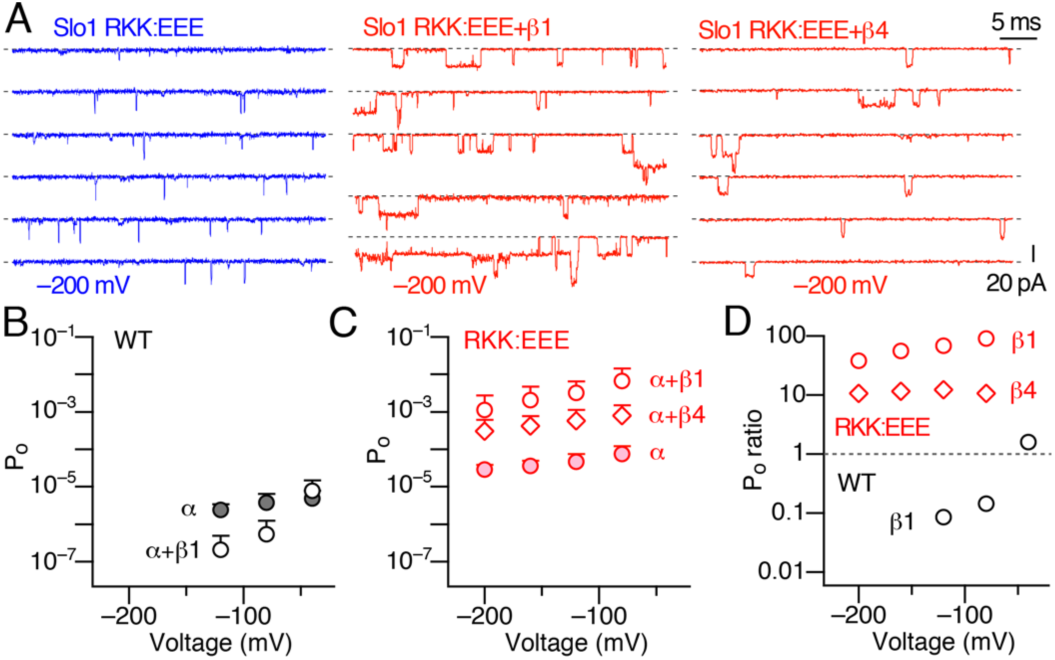
Single-channel openings at extreme negative voltages without Ca^2+^. **A**. Illustrative openings at –200 mV from hSlo1 RKK:EEE (*left*), hSlo1 RKK:EEE+β1 (*center*), and hSlo1 RKK:EEE+β4 (*right*). The membrane patches shown contained about 40, 34, and 26 channels, respectively. Dashed lines indicate zero-current levels. **B**. Voltage dependence of P_o_ in wild-type hSlo1 (filled circles) and hSlo1+β1 (open circles). n = 14. **C**. Voltage dependence of P_o_ in hSlo1 RKK:EEE (filled circles), hSlo1 RKK:EEE+β1 (open circles), and hSlo1 RKK:EEE+β4 (open diamonds). n = 8 - 10. **D**. Fractional increases in P_o_ by coassembly of hSlo1 RKK:EEE with β1 (red circles) and with β4 (red diamonds). The results for wild-type hSlo1+β1 are also shown with black circles. n = 6 - 12.

### hSlo1 RKK:EEE+β1 and hSlo1 RKK:EEE+β4 remain Ca^2+^ sensitive

Changes in L_0_ such as those by inclusion of β subunits are not expected to markedly impair the overall Ca^2+^ sensitivity, changes in V_0.5_ by Ca^2+^, of the resulting channels. Consistent with this prediction, hSlo1 RKK:EEE+β1 and hSlo1 RKK:EEE+β4 remained sensitive to intracellular Ca^2+^ (Fig. S6). The change in V_0.5_ by increasing [Ca^2+^]_i_ from 0 to 100 µM in hSlo1 RKK:EEE+β1, about –350 mV (Fig. S6 circles), was similar to that observed with wild-type hSlo1+β1 (9). The similar shifts in V_0.5_ observed in wild-type hSlo1+β1 and hSlo1 RKK:EEE+β1 by Ca^2+^ may also argue against the idea that hSlo1 RKK:EEE+β1 with a left-shifted V_0.5_ value without Ca^2+^ somehow resembles Ca^2+^-bound hSlo1+β1. The GV curve with 100 µM [Ca^2+^]_i_ is consistent with the HA model prediction (Fig. S6) using a greater value of L_0_ for hSlo1 RKK:EEE (20) and the adjustments made by Orio & Latorre for inclusion of β1 (9). Additionally, the change in V_0.5_ observed with hSlo1 RKK:EEE+β4 by increasing [Ca^2+^]_i_ to 100 µM (Fig. S6 diamonds) was also comparable to that in wild-type Slo1+β4 (10).

### Coassembly with γ1 (LRRC26)

A large negative shift in V_0.5_ is also observed when hSlo1 coassembles with γ1 (LRRC26) (12). The underlying mechanism of the γ1-mediated shift is a drastic increase in the strength of gate-voltage sensor coupling as specified by the parameter D in the HA model without a noticeable change in L_0_ (12) (Fig. S7 *top*). Coexpression of hSlo1 RKK:EEE with γ1 moved the resulting complex’s V_0.5_ to the negative direction by a magnitude similar to that found with wild-type hSlo1 (Fig. S7 *bottom*). The action of γ1 is unlikely to depend on the RKK segment in the same way that that of β1/β4 does.

### Loci responsible for the greater effect of β1 compared with β4

The intracellular N terminus of *β* subunits is involved in regulation of the equilibrium constant L_0_ by select free fatty acids (16). Prompted by this previous finding, mutations in the β1 N terminus were made to investigate why β1 induces a greater shift in V_0.5_ than β4 when coassembled with hSlo1 RKK:EEE (Fig. 1). Sequence comparison of the N-terminal regions of β1 and β4 reveals one noticeably divergent area between the two subunits: ^8^AQKR^11^ in *β*1 and ^9^EYTE^12^ in β4 (Fig. 3A); the segment in *β*1 appears to more positive than that in β4. The residues in this segment were exchanged between β1 and β4 (Fig. 3B). The GV curve from hSlo1 RKK:EEE+β1 A8E:Q9Y:K10T:R11E (hereafter referred to as AQKR:EYTE), inserting β4 ^9^EYTE^12^ into the β1 background, was essentially indistinguishable from that of hSlo1 RKK:EEE+ *β*4 (Fig. 3C *first row* blue circles). Conversely and importantly, the GV curve from hSlo1 RKK:EEE+*β*4 E9A:Y10Q:T11K:E12R (EYTE:AQKR), with β1 ^8^AQKR^11^ residues in the *β*4 background, closely resembled that of hSlo1 RKK:EEE+β1 (Fig. 3C *first row* black diamonds). The residues ^8^AQKR^11^ in β1 and ^9^EYTE^12^ in β4 thus largely account for the V_0.5_ difference between hSlo1 RKK:EEE+β1 and hSlo1 RKK:EEE+β4. The same residues also explain the differences in current relaxation time course at > 0 mV (Fig. 3D *first row*). However, we note that the kinetics at more negative voltages of hSlo1 RKK:EEE+β1 AQKR:EYTE and hSlo1 RKK:EEE+β4 EYTE:AQKR did not exactly match those of hSlo1 RKK:EEE+β4 or hSlo1 RKK:EEE+β1 (Fig. 3D *first row*). The mutations β1 K10Y:R11E (KR:TE) and β4 T11K:E12R (TE:KR), exchanging only two of the four residues in the aforementioned β segment (Fig. 3A), were less effective in rendering the GV and current kinetics of the resulting channels to resemble those of hSlo1 RKK:EEE+β4 and hSlo1 RKK:EEE+β1, respectively (Fig. 3C-D *second row*). The single-residue exchange mutations β1 R11E/β4 E12R and β1 A8E/β4 E9A were even less effective (Fig. 3C-D *third and fourth rows*).

**Figure 3.**
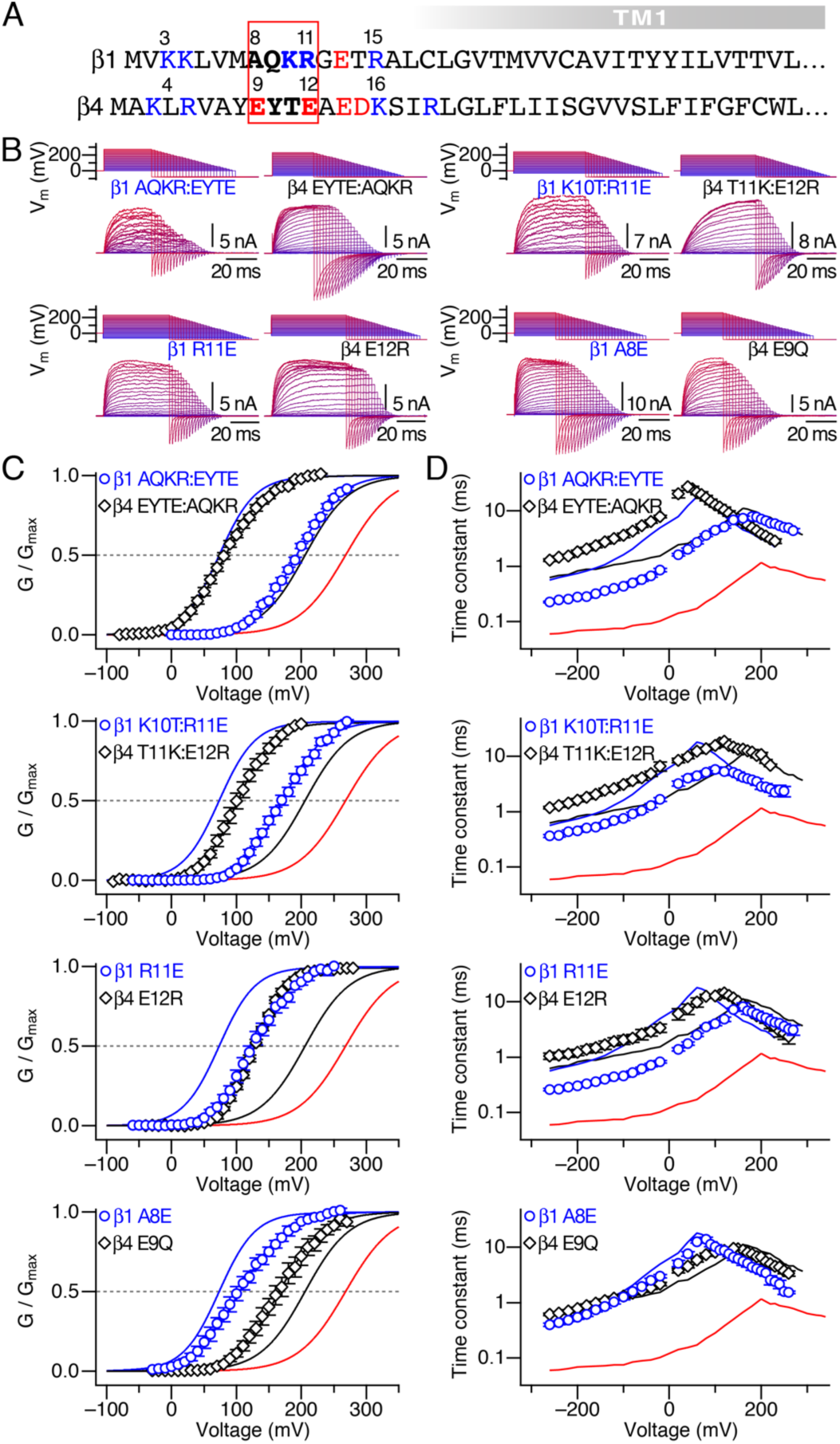
N-terminal exchange mutations in β1 and β4. **A**. N-terminal sequences of β1 and β4. **B**. Illustrative currents from hSlo1 complexes containing β1 with β1-to-β4 mutations and those containing β4 with β4-to-β1 mutations. **C**. Normalized GV curves from the currents shown in **B**. See Table S5 for GV parameters. **D**. Time constants of current relaxation from the results in **B**. In each graph in **C** and **D**, the red, blue, and black solid curves represent the results from hSlo1 RKK:EEE, hSlo1 RKK:EEE+β1, and hSlo1 RKK:EEE+β4, respectively, for comparison.

E13 and T14 in β1, C-terminal to ^8^AQKR^11^, have been suggested to interact with the residues near the Slo1 RCK1 Ca^2+^ sensor (Fig. 1A) in the cytoplasmic gating ring (GR) domain electrostatically to regulate Ca^2+^-dependent gating of the Slo1+β1 channel (18). Neutralization of E14, D15, and K16 in β4 to Gln, however, had only a modest impact on the overall voltage dependence of activation and current relaxation without Ca^2+^ (Fig. S8).

### Interactions between hSlo1 ^329^RKK^331^ and β1 ^8^AQKR^11^

To determine whether the hSlo1 mutation RKK:EEE and the β1 ^8^AQKR^11^ mutations function in an independent or a non-independent fashion, double mutant cycle analysis (23, 30) was performed. The values of the equilibrium constant L_0_ in the single and double mutants were estimated from P_o_ measurements at extreme negative voltages without Ca^2+^. The β1 ^8^AQKR^11^ segment was mutated to single amino-acid repeats (e.g., EEEE) to facilitate data interpretation. Both hSlo1 RKK:EEE and β1 AQKR:EEEE, especially the former, individually increased the value of L_0_, leading to a net stabilization of the closed conformation (Fig. 4A). The greater P_o_ value in hSlo1 RKK:EEE+β1 is immediately obvious (Fig. 4A *top right*). However, the P_o_ value in the double mutant hSlo1 RKK:EEE+β1 AQKR:EEEE is clearly smaller than that in hSlo1 RKK:EEE+β1 (Fig. 4A *bottom right*), revealing a non-independent action of the two mutations hSlo1 RKK:EEE and β1 AQKR:EEEE. Accordingly, the degree of coupling between the hSlo1 mutation RKK:EEE and the β1 mutation AQKR:EEEE revealed by the mutant cycle analysis, 4.3 x 10^−3^, markedly deviated from unity, indicative of a strong interaction between the two mutations. The coupling factor values estimated for other β1 mutants based on the macroscopic HA model fitting process (see Materials and Methods) generally deviated from unity and ranged from 8 in AQKR:RRRR to 4.3 x 10^−3^ in AQKR:EEEE (Fig. 4B; Fig. S9), suggesting that the hSlo1 mutation RKK:EEE and select β1 AQKR mutations can exert non-independent effects on L_0_ and thus GV V_0.5_. The only exception was β1 AQKR:GGGG; the coupling factor value of ∼1 for hSlo1 RKK:EEE and β1 AQKR:GGGG indicates that the β1 side-chain moieties may be important.

**Figure 4.**
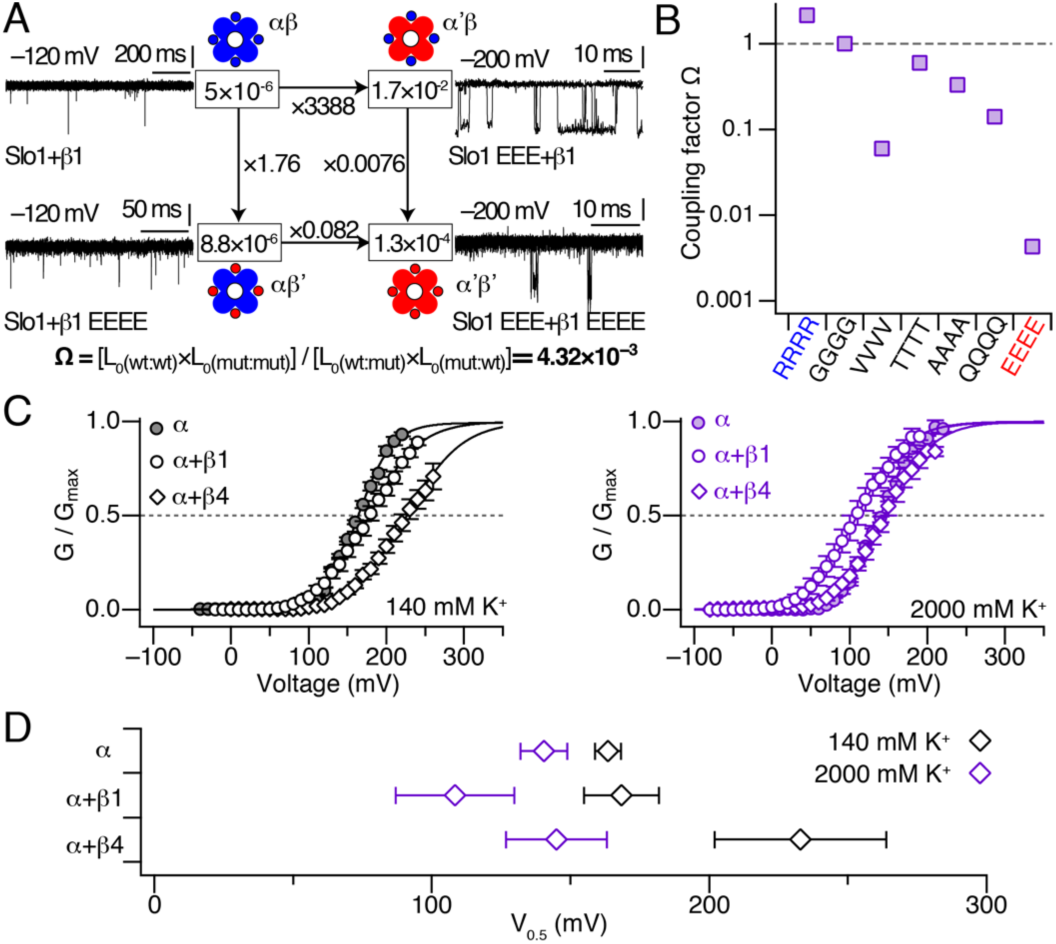
Electrostatic interactions between hSlo1 ^329^RKK^331^ and β1 ^8^AQKR^11^. **A**. Double-mutant cycle analysis using single-channel openings at negative voltages without Ca^2+^. The two α subunits used were WT hSlo1 (“α” in this figure) and hSlo1 RKK:EEE (“EEE”; “α’” in this figure). The β1 subunits used were WT β1 (“β” in this figure) and β1 ^8^AQKR^11^: ^8^EEEE^11^ (“EEEE”; “β1EEEE*”* in this figure). The recording voltages were at least 120 mV more negative than the respective GV V_0.5_ values and indicated. Ten data sweeps are superimposed. The L_0_ values estimated from the single-channel P_o_ values (see Materials and Methods) are indicated in the boxes and fractional changes in L_0_ are indicated along the arrows connecting the L_0_ boxes. All vertical scale bars represent 10 pA. **B**. Coupling factor values with different β1 mutants in which ^8^AQKR^11^ are replaced with the residues indicated. The two α subunits used were WT hSlo1 and hSlo1 RKK:EEE. The coupling factor values were calculated from the L_0_ values estimated from fitting macroscopic currents with the HA model (see Materials and Methods). See Table S6 for GV parameters. **C**. Normalized GV curves of hSlo1 (filled circles), hSlo1+β1 (open circles), and hSlo1+β4 (open diamonds) with 140 mM KCl (*left*) and 2000 mM KCl (*right*). Each patch was tested under the two conditions. For GV parameter values, see Table S7. **D**. Changes in GV V_0.5_ by increasing [KCl] from 140 mM (black diamonds) to 2000 mM (purple diamonds) in hSlo1, hSlo1+*β*1, and hSlo1+*β*4. n = 7 to 10.

### Electrostatic interactions between wild-type hSlo1 and *β*1

To infer whether wild-type hSlo1 with ^329^RKK^331^ and wild-type β1 with ^8^AQKR^11^ or with wild-type β4 with ^9^EYTE^12^ interact electrostatically, electrophysiological measurements from hSlo1, hSlo1+β1, and hSlo1+β4 were made under two different concentrations of KCl in the intracellular side: 140 mM and 2000 mM. For these KCl concentrations, the calculated Debye lengths are about 0.8 and 0.2 nm, respectively, assuming the relative permittivity value of 80. In comparison, the RKK-EE ring stabilizing the closed conformation has a diameter of about 4 to 5 nm (Fig. 5). With 140 mM KCl, inclusion of β subunits, especially of β4, shifts V_0.5_ to the positive direction (Fig. 4C *left*). With 2000 mM KCl, the V_0.5_ shift by β4 was absent and β1 induced a small negative V_0.5_ shift (Fig. 4C *right*, D). The V_0.5_ values of hSlo1, hSlo1+β1, and hSlo1+β4 with 140 and 2000 mM KCl are summarized in Fig. 4D.

**Figure 5.**
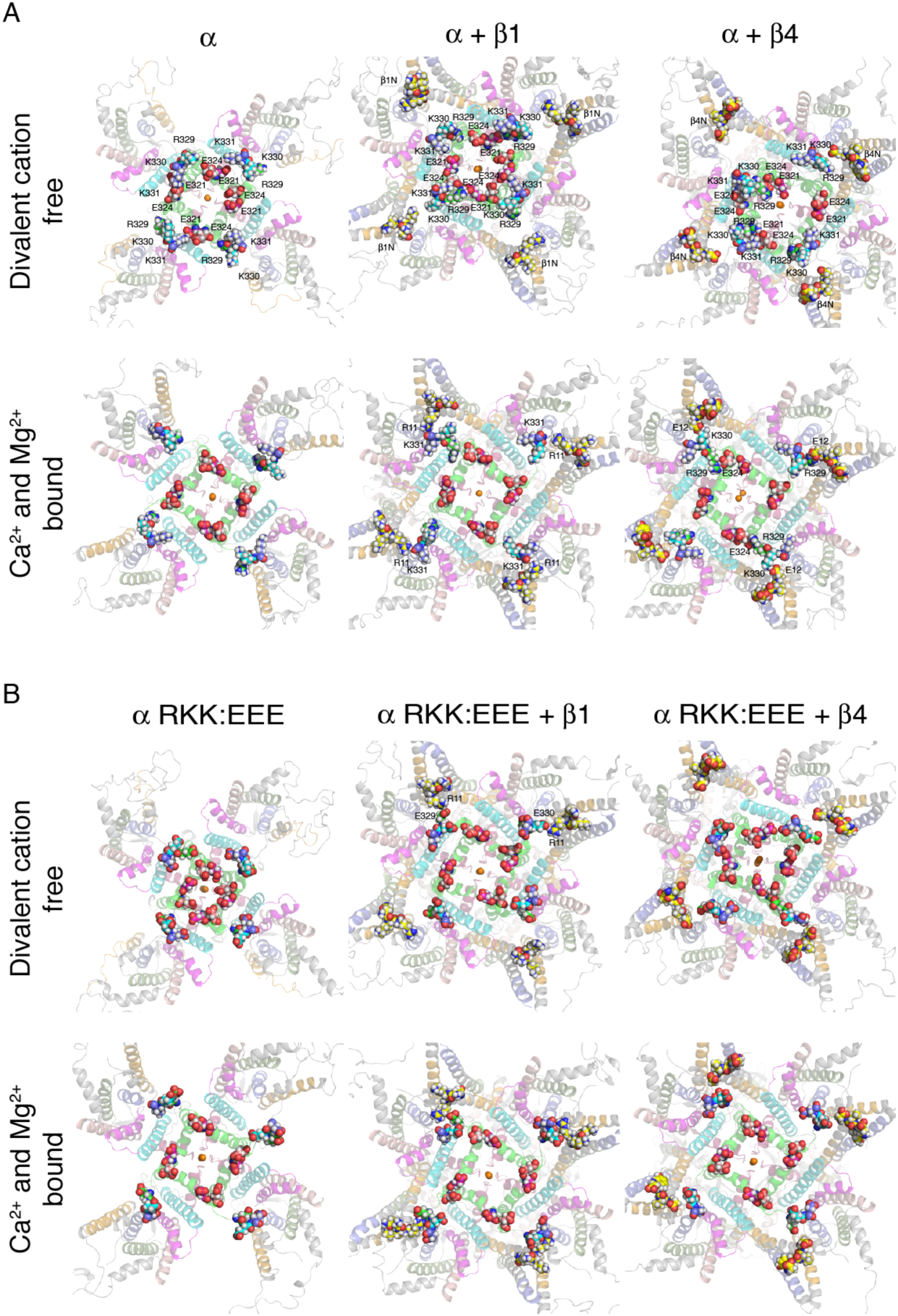
Snapshots of the structures optimized by MD simulation runs. **A**. Wild-type hSlo1 alone, hSlo1+β1, and hSlo1+β4. **B**. hSlo1 RKK:EEE, hSlo1 RKK:EEE+β1, and hSlo1 RKK:EEE+β4. Each channel type was studied without divalent cations *(top row* in each panel) and with Ca^2+^ and Mg^2+^ bound (*bottom row* in each panel). The structures are viewed from the intracellular side. The intracellular domain of the channel C-terminal to the RKK residues is not shown. Water, lipids, and ions other than the pore K^+^ ions are not shown. Different residues are rendered using different carbon-color schemes; E321: magenta, E324: salmon, R329/E329: green, K330/E330: cyan, K331/E331: blue, β1 AQKR and β4 EYTE: yellow.

One major confounding factor in the ionic-strength approach is that such manipulations have potential to alter multiple aspects of hSlo1 gating. Our mutant cycle results suggest that the mutations hSlo1 RKK:EEE and β1 AQKR:GGGG appear to function in an independent fashion (see Fig. 4B). Increasing [KCl] from 140 to 2000 mM in hSlo1+β1 AQKR:GGGG only modestly changed its V_0.5_ (Fig. S10). The shift in V_0.5_ was similar to that observed with hSlo1 alone (Fig. 4D), suggesting that the greater shifts in V_0.5_ observed in hSlo1+β1, and hSlo1+β4 (Fig. 4D) with high KCl may involve β1 ^8^AQKR^11^ and β4 ^9^EYTE^12^.

### Inferences from hSlo1 structural models

Structural models of hSlo1 (13) and hSlo1 RKK:EEE (see Materials and Methods), with and without β1/β4, in membranes were optimized by molecular dynamics (MD) simulations to infer the structural differences around the ^329^RKK^331^ segment. Consistent with our previous simulation results (20), ^329^RKK^331^ and E321/E324 were observed to form the RKK-EE ring in wild-type hSlo1 without divalent cations (Fig. 5A left*, top row*). Without Ca^2+^ or Mg^2+^, the overall open probability value of the wild-type hSlo1 channel at 0 mV is < 1 x 10^−4^ (4) and this structure thus probably represents a closed conformation. In wild-type hSlo1, with Ca^2+^ and Mg^2+^ fully bound, the open probability is > 0.5 at 0 mV (4) and the RKK-EE ring is impaired (Fig. 5A, *left, second row*); the RKK residues moves away from the pore axis and stabilized by nearby membrane lipids as shown previously (20). In the presence of β1 or β4, when Ca^2+^ and Mg^2+^ are bound, the RKK residues generally move away from the pore axis and approach the N-terminal residues of β1/4 (Fig. 5A, *middle* and *right* columns, *second* rows). In the presence of β1 with Ca^2+^ and Mg^2+^ bound, the backbone oxygen atoms of K330 and K331 interact with the side chains of β1 R11 {Hoshi, 2013 #5819}. Only occasionally, the side chain of R329 interacts with E324. In the presence of β4 and Ca^2+^ and Mg^2+^ bound, β4 E12 interact with R329 as in the original cryo-EM structure and also with K330 (Fig. 5A, *right*). We note that when K330 of one subunit is interacting with β4 E12, the side chain of R329 of the same subunit is often in contact with that of E324.

In hSlo1 RKK:EEE, the four sets of E321/E324 appear to form a smaller ring (Fig. 5B, *left*, *third* row;) than in wild-type hSlo1 (Fig. 5A, *left*, *first* row) without divalent cations, presumably because of the repulsive electrostatic interactions between ^329^EEE^331^ and E321/. In hSlo1 RKK:EEE+β1 and hSlo1 RKK:EEE+β4 without divalent cations, some of the RKK:EEE segments interact with the β1/β4 N-terminal residues (Fig. 5B *middle* & *right*, *third* row). The overall arrangements of RKK:EEE and E321/E324 in these channels without divalent cations resemble those of wild-type hSlo1+β1 and hSlo1+β4 with Ca^2+^ and Mg^2+^ bound. The Ca^2+^ and Mg^2+^-bound structures of the channel complexes are not noticeably altered by the triple mutation RKK:EEE (Fig. 5B *second* vs. *fourth* rows).

As noted earlier (see Fig. 4), the presence of 2000 mM KCl diminishes the negative-going shift in V_0.5_ by coassembly with β1 or β4 in wild-type hSlo1. In wild-type hSlo1+β1 optimized with 2000 mM KCl, ^329^RKK^331^ residues break away from E321/E324 (Fig. S16), somewhat like the arrangement seen with hSlo1 with Ca^2+^ and Mg^2+^ bound (see Fig. 5). The oxygen atoms in E321/E324 interact with K^+^ ions (Fig. S16 orange spheres) and Cl^−^ ions (Fig. S16 green spheres) are found near the RKK segments as well as *β*1 N terminus (Fig. S16 green spheres).

## Discussion

Our macroscopic current measurements presented here show that coassembly of hSlo1 with auxiliary β1 or β4 subunits moves the overall voltage dependence of the channel complex without Ca^2+^ to the negative direction by as much as 190 mV, depending on the amino acids at positions 329-331 in hSlo1 immediately C-terminal to S6. Further, the single-channel measurements demonstrate that the aforementioned shifts in the overall voltage dependence are caused primarily by alterations in the intrinsic behavior of the ion conduction gate without allosteric influences from the voltage or divalent cation sensors as specified by the equilibrium constant L_0_ in the HA model of Slo1 gating (4). The results of the double-mutant cycle analysis and ionic-strength experiments further suggest that the residues at positions 329-331 in hSlo1 are capable of forming electrostatic interactions with those at β1 positions 8-11 and β4 positions 9-12 facing the intracellular side.

In our previous study (20) we showed that the closed conformation of the ion conduction gate is stabilized in part by inter-subunit ion-ion interactions between ^329^RKK^331^ and E321/E324 in S6 and ^329^RKK^331^. The open conformation of the gate, in contrast, is stabilized partly by the ion-dipole interactions between ^329^RKK^331^ and the oxygen atoms of nearby membrane lipids (20). A similar lipid interaction has been also suggested by Yazdani et al. (21). The above framework may be used to account for the observations presented in this study.

In this study, the energy-minimized structures of hSlo1, hSlo1+β1, hSlo1+β4, hSlo1 RKK:EEE, hSlo1 RKK:EEE+β 1, and hSlo1 RKK:EEE+β4 were estimated using MD simulations under two conditions: divalent cation free and Ca^2+^ and Mg^2+^ bound. The main features of the hSlo1 structures presented here are indistinguishable from those of our earlier study (20), increasing the level of credence in the simulation results.

Coassembly of wild-type hSlo1 with β1 or β4 leads to a net stabilization of the closed conformation of the gate, decreasing L_0_ (9, 10). However, both the closed and open conformations are more stable than in wild-type hSlo1 because the activation and deactivation kinetics are noticeably slower with β1 or β4. Inspection of the energy-minimized structures of hSlo1, hSlo1+β1, and hSlo1+β4 without divalent cations, mostly likely representing closed conformations, show that the RKK-E321/E324 interactions remain intact, consistent with the importance of the RKK-EE ring in closed-state stabilization. However, it is unclear whether the RKK-EE interactions in hSlo1+β1 and hSlo1+β4 structures differ appreciably from those in hSlo1 to account for the slower activation kinetics in hSlo1+β1 and hSlo1+β4. In the Ca^2+^- and Mg^2+^-bound structures, presumably representing open conformations, the RKK residues interact with β1/4 N-terminal residues. With β1, the RKK residues interact with the backbone oxygen atoms of *β*1 R11. With β4, the RKK residues interact with side-chain oxygen atoms of β4 E12. These protein-protein interactions in hSlo1+β1 and hSlo1+β4 must be more effective than the RKK-lipid interactions observed in hSlo1 (20) in open-state stabilization; the deactivation kinetics are slower with β1 or β4.

The energy-minimized structures of the hSlo1 RKK:EEE channels offer clues as to why coassembly with β1 or β4 increases L_0_ and produces a negative-going shift in V_0.5_. In hSlo1 RKK:EEE without divalent cations, the repulsive ion-ion interactions between ^329^EEE^331^ and E321/E324 disrupt both the closed-state stabilizing RKK-EE interactions and the open-state stabilizing RKK-lipid interactions. The former is predominant and increases the L_0_ value. With β1 present, ^329^EEE^331^ residues now expand outward from the pore axis towards the *β*1 N termini (^8^AQKR^11^), forming ion-ion interactions with *β*1 R11; this arrangement between ^329^EEE^331^ and the β1 N terminus is reminiscent of the wild-type ^329^RKK^331^-β1 structural arrangement with Ca^2+^ and Mg^2+^ bound (Fig. 5). We thus postulate that the expanded ^329^EEE^331^ arrangement reaching towards β1 R11 contributes to the increased L_0_ and thus the leftward shift in GV V_0.5_. The N terminus of *β*4 does not contain a residue equivalent to β1 R11 but instead possess two negatively charged residues (^9^EYTE^12^). ^329^EEE^331^, E321/E324, and the β4 N termini primarily interact with water and K^+^ and this arrangement does not increase L_0_ as much as that found with β1. When Ca^2+^ and Mg^2+^ ions are bound, ^329^EEE^331^ are stabilized by water and K^+^. The above scenario suggests that coassembly of hSlo1 RKK:EEE with β1 destabilizes the closed state more effectively or increasing the gate opening rate constant compared with wild-type hSlo1+β1. This postulate is consistent with the ∼8 to 10-fold faster activation kinetics of hSlo1 RKK:EEE+*β*1 compared with wild-type hSlo1+β1 at positive voltages (Fig. 1E). The deactivation kinetics of hSlo1 RKK:EEE+β1 is two to three times slower than that in wild-type hSlo1+β1. The slower deactivation could be accounted for the interactions of the side-chain negative charges of ^329^EEE^331^ with the side chain positive charge of *β*1 R11. We note that the deactivation kinetics of hSlo1 RKK:EEE+β1 and hSlo1 RKK:EEE+β4 are similar (Fig. 1E). In the Ca^2+^- and Mg^2+^- bound structure of hSlo1 RKK:EEE+ *β*4, ^329^EEE^331^ residues interact with water and K^+^. It is perplexing that the interactions between ^329^EEE^331^ and β1 R11 in hSlo1 RKK:EEE+β1 and those between ^329^EEE^331^ with water and K^+^ in hSlo1 RKK:EEE+β4 are similar in strength to produce the near identical deactivation kinetics.

The results of the mutant cycle analysis covering the RKK and β1 ^9^EYTE^12^ segments are generally consistent with the features in the energy-minimized structures discussed above. For example, the finding that the mutation RKK:EEE and the β1 mutation EYTE:EEEE do not function independently on the equilibrium constant L_0_ is easily understood in terms of the electrostatic repulsion forces separating ^329^EEE^331^ and β1 EEEE (see Fig. 5). Further, the finding that 2000 mM KCl greatly diminished the effect of coassembly of wild-type hSlo1 with β1 on L_0_ and thus V_0.5_ (Fig. 4) is also readily understood; the structural simulation shows preferential localization of K^+^ ions near E321 and E324 and that of Cl^−^ ions near ^329^RKK^331^ and β1 N terminus, acting to screen the electrostatic interactions among ^329^RKK^331^, E321/E324, and β1 N terminus (Fig. S16).

The results presented here focusing on L_0_ in the absence of Ca^2+^ may be compared with those of an earlier study by Liu et al. performed with ≥ 10 µM Ca^2+^ (18). Using macroscopic GV curves, Liu et al. inferred the free energy changes associated with changes in the apparent voltage dependence of mouse Slo1+β1 with Ca^2+^. The analysis suggested that β1 ^13^ET^14^, just C-terminal to β1 ^8^AQKR^11^ examined in this study, interacts with the hSlo1 ^392^KR^393^-equivalent residues in the RCK1 area (see Fig. 1A) in the GR domain. The interaction between hSlo1 ^392^KR^393^ and β1 ^13^ET^14^ reported by Liu et al. is unlikely to contribute to the results presented here obtained without Ca^2+^. Neutralization of β1 ^13^ETR^15^ to Gln essentially has no impact (Fig. S8). Further, coassembly with β1 increases open probability in the hSlo1 RKK:EEE channel lacking the cytoplasmic GR domain including the RCK1 area (31, 32) (Fig. S11).

The framework focusing on ^329^RKK^331^, E321/E324, and β1/4 N terminus qualitatively describes the manner by which the said components regulate the overall stability of the ion conduction gate or the equilibrium constant L_0_. Clearly further work will be need to assess how ^329^RKK^331^, E321/E324, and β1 N terminus energetically and quantitatively contribute to the opening and closing rate constants of the ion conduction gate. Undoubtedly, such calculations will require careful consideration of the contributions from ions, water and lipids.

Our study illustrates the importance of ^329^RKK^331^ and E321/E324 in regulation of a specific step in gating of Slo1 channels, the ion conduction gate specified by the equilibrium constant L_0_, without or with auxiliary β subunits. This constant L_0_ is subject to regulation by a variety of modulators (33), including DHA (34). While the action of DHA involves an ion-dipole interaction between the carboxylic acid group of DHA and the OH group of hSlo1 Y318 in S6, it is possible that the consequence of this interaction is somehow transmitted to the final effector, the ion conduction gate, through ^329^RKK^331^ and E321/E324. Similarly, other hydrophobic modulators including lipids and cholesterols (35) may impart their action through ^329^RKK^331^.

Exactly how the interactions involving ^329^RKK^331^, E321/E324, and β1/4 N terminus lead to changes in the ion conduction gate machinery located elsewhere, most probably near the ion selectivity filter, requires further investigation. The detailed nature of the Slo1 ion conduction gate has remained largely elusive. While it is agreed that the S6 helices do not form a tight hydrophobic seal and that the selectivity filter may be important (36-38), a clear physicochemical view of the ion conduction gate is yet to emerge. Functionally, the gate behavior is known to be altered by various mutations, especially those in S6 (39), and by many small molecules including “BK openers” with potential therapeutic potential (33). Alterations of the RKK-EE ring may represent an important way for small molecules to regulate the behavior of the ion conduction gate of Slo1. Schewe et al., examining multiple channel types, have revealed that the region just intracellular to the selectivity filter segment is a critical locus for drug binding in Slo1 and other channels (40). It is plausible that the allosteric influence from the RKK and E321/E324 segments converge onto the selectivity gate through the drug binding motif identified by Schewe et al.

In summary, this study demonstrates that the ion conduction gate of the hSlo1 channel is critically regulated by ^329^RKK^331^ and E321/E324. This functional regulation is observed in hSlo1 channels as well as in hSlo1 complexed with β1 or β4 through electrostatic interactions. The ^329^RKK^331^ and E321/E324 may be involved in the mechanisms by which small modulators of the channel influence the ion conduction gate.

## Acknowledgements

The study was supported in part through the National Science Foundation of China (YT NSCF 32071103), NIH (TH R01GM121375) and the German Research Foundation (SHH HE2993/18-1).

---

**Figure S1.**
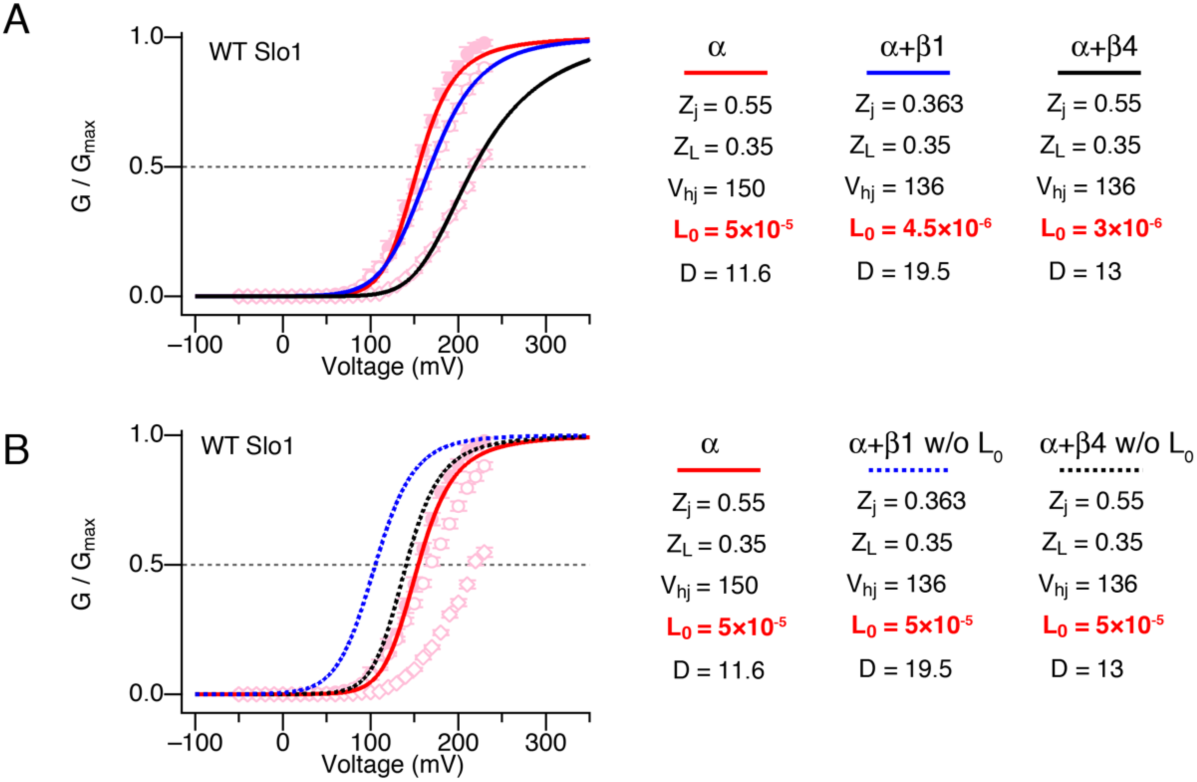
GV changes by β1/4 explained by the HA model. **A**. Normalized GV curves from hSlo1, hSlo1+ β1, and hSlo1+β4 fitted by the HA model. The HA model parameters were changed according to the adjustments made by Orio and Latorre (9) and the L_0_ values were fine-tuned using the single-channel results at negative voltages (see Materials and Methods). The parameter values are shown right of the graph. **B**. HA model fits without changing the L_0_ value.

**Figure S2.**
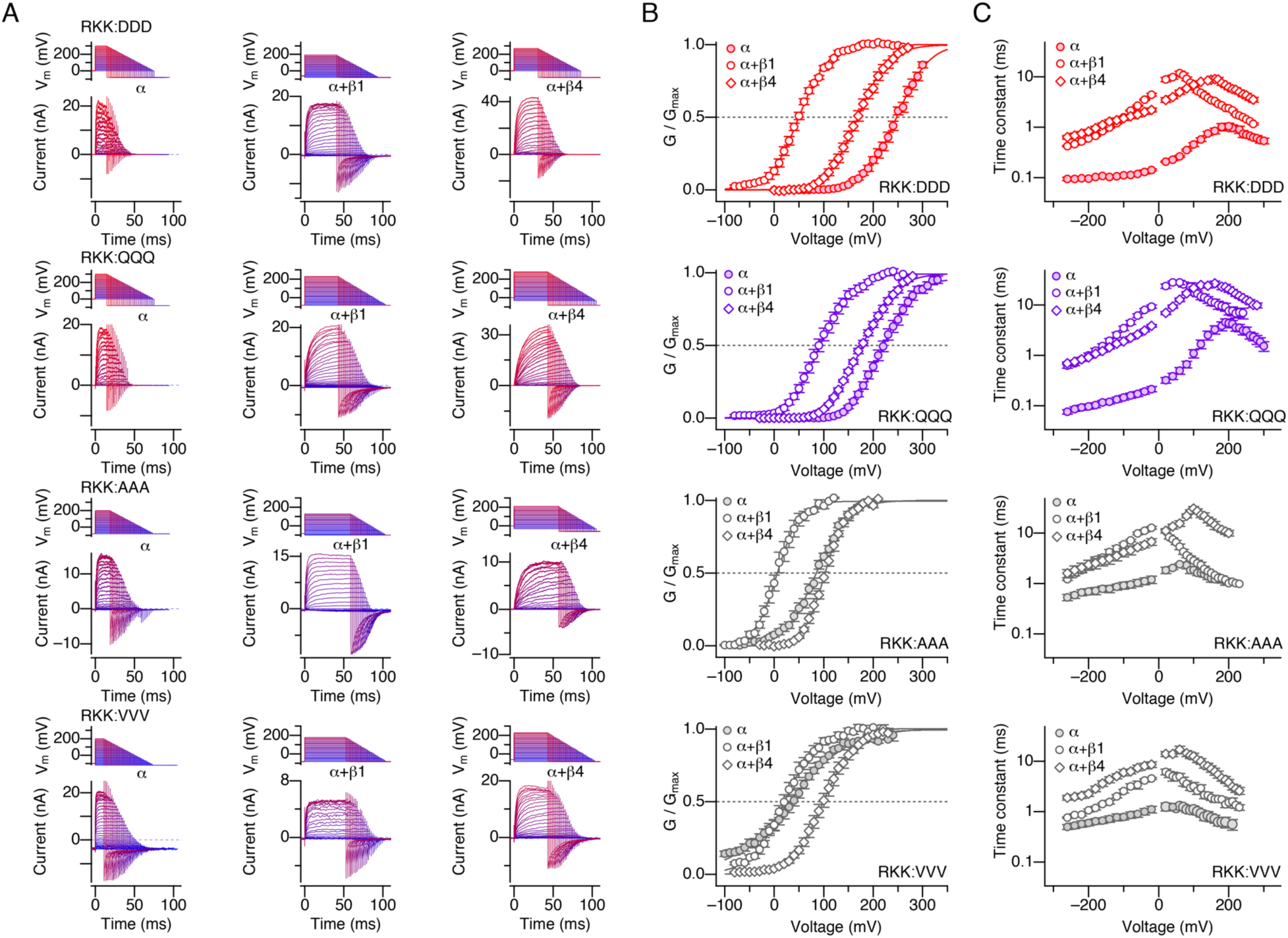
hSlo1 RKK mutants coassembled with β1 or β4. **A**. Illustrative currents from α alone (*left*), with β1 (*center*), and with β4 (*right*) of the hSlo1 RKK mutants indicated. From *top* to *bottom*: RKK:DDD, RKK:QQQ, RKK:AAA, and RKK:VVV. The voltage and current sweeps use the same color scheme, from hyperpolarized (blue) to depolarized (red) test voltages. **B**. Normalized GV curves from α alone (filled circles), with β1 (open circles), and with β4 (open diamonds) of the mutants shown in **A**. Smooth curves are Boltzmann-type fits (see Table S1). **C**. Time constants of current relaxation at different voltages from α alone (filled circles), with β1 (open circles), and with *β*4 (open diamonds). All results were obtained without Ca^2+^.

**Figure S3.**
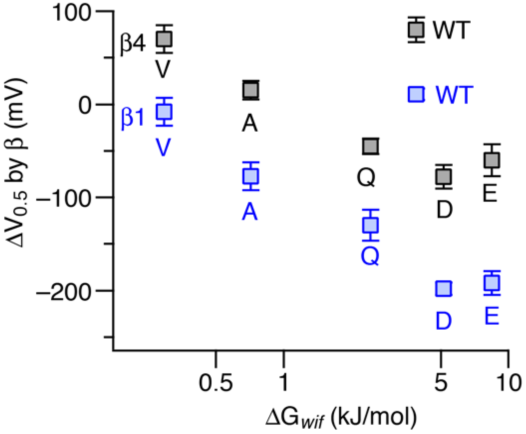
Changes in V_0.5_ by β1 or β4 as a function of the Wimley-White hydrophobicity (ΔG*_wif_*) index of the amino acid in the RKK segment in the mutants in which each of the ^329^RKK^331^ residues is substituted with the amino acid indicated. The channels contained β1 (blue) or β4 (black). The V_0.5_ results are relative to those without β (Table S1). All results were obtained without Ca^2+^.

**Figure S4.**
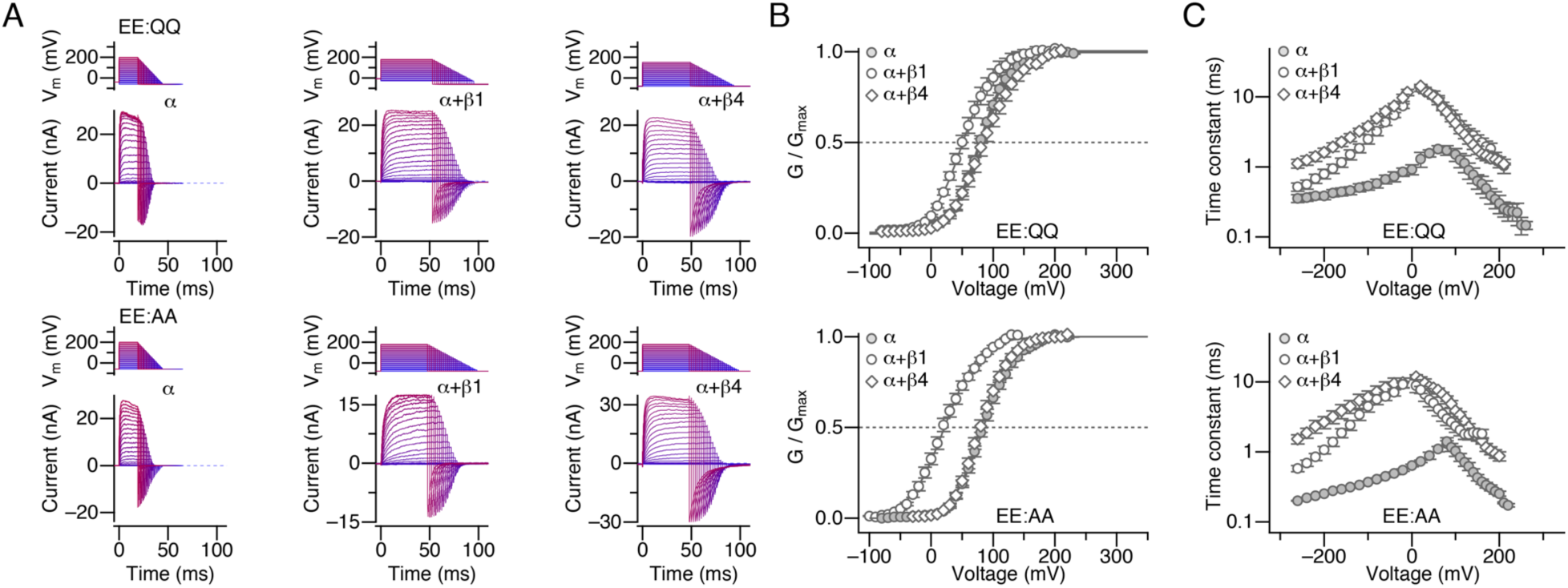
Changes in currents through hSlo1 E321Q:E324Q (EE:QQ) (*top*) and hSlo1 E321A:E324A (EE:AA; *bottom*) by β1 and β4. **A**. Illustrative currents through hSlo1 alone (*left*), with β1 (*center*), and with β4 (*right*). **B**. Normalized GV curves from α alone (filled circles), with β1 (open circles), and with β4 (open diamonds). Smooth curves are Boltzmann-type fits (see Table S2). **C**. Time constants of ionic current relaxation at different voltages from α alone (filled circles), with β1 (open circles), and with β4 (open diamonds). All results were obtained without Ca^2+^.

**Figure S5.**
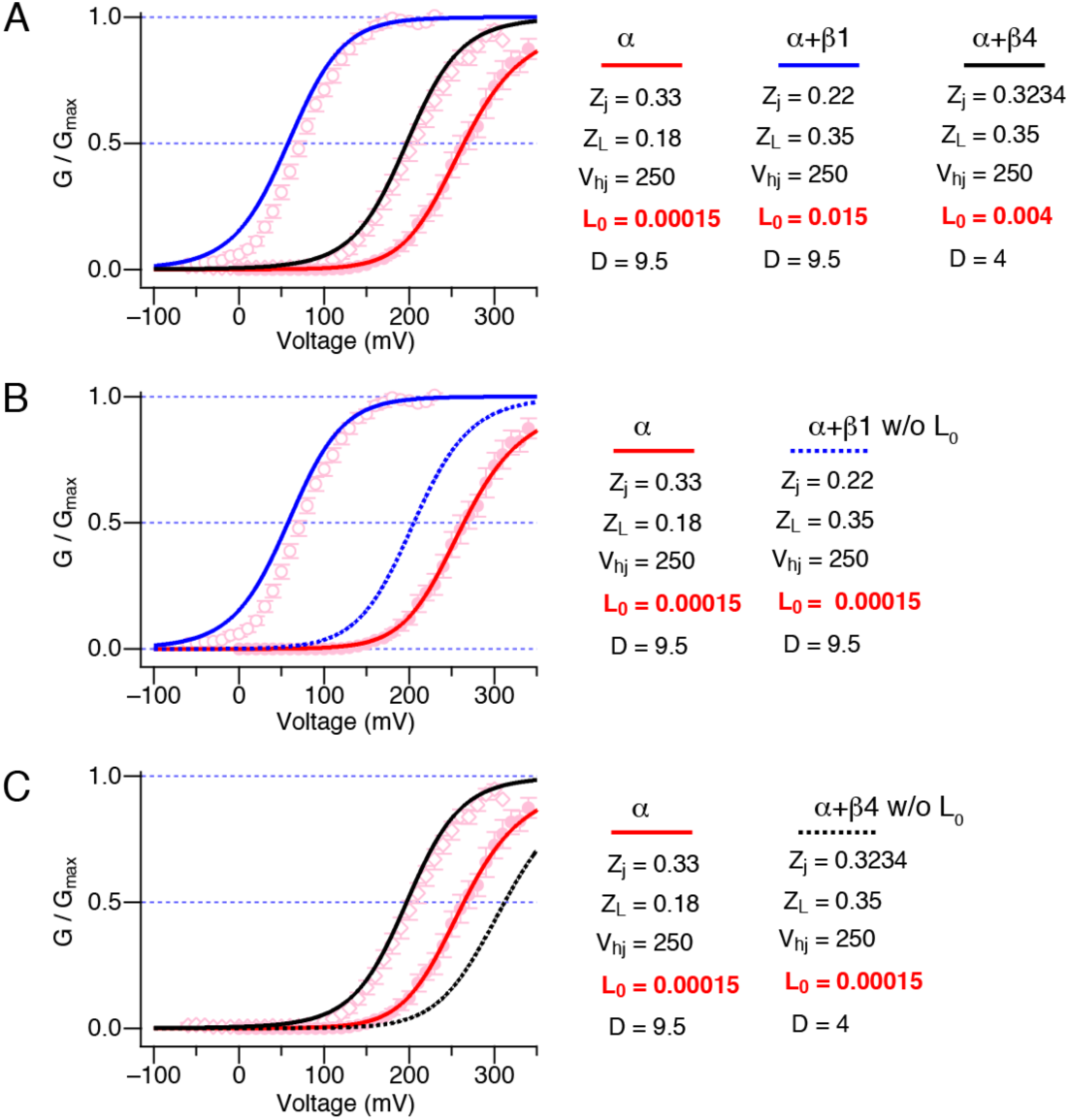
Changes in L_0_ explain GV changes of hSlo1 RKK:EEE by β1/ β4. **A**. Normalized GV curves from hSlo1 RKK:EEE (filled circles), hSlo1 RKK:EEE+β1 (open circles), and hSlo1 RKK:EEE+β4 (open diamonds). The smooth solid curves (red: hSlo1, blue: hSlo1+β1, black: hSlo1+β4) are the predictions of the HA model using the parameter values shown (*right*). The L_0_ values were determined directly from the single-channel measurements at extreme negative voltages without Ca^2+^ (see Materials and Methods). **B**. Normalized GV curves from hSlo1 RKK:EEE (filled circles), hSlo1 RKK:EEE+β1 (open circles). The solid smooth curves the HA model predictions as in **A**. (red: hSlo1, blue: hSlo1+β1). The blue dashed curve shows the HA model prediction without changing the L_0_ value. **C**. Normalized GV curves from hSlo1 RKK:EEE (filled circles) and hSlo1 RKK:EEE+β4 (open circles). The solid smooth curves (red: hSlo1, black: hSlo1+β4) are the predictions of the HA model as in **A**. The black dashed curve shows the HA model prediction without changing the L_0_ value. Notice that the predicted GV curves with L_0_ constrained deviate markedly from the experimentally observed curves, illustrating the importance of L_0_.

**Figure S6.**
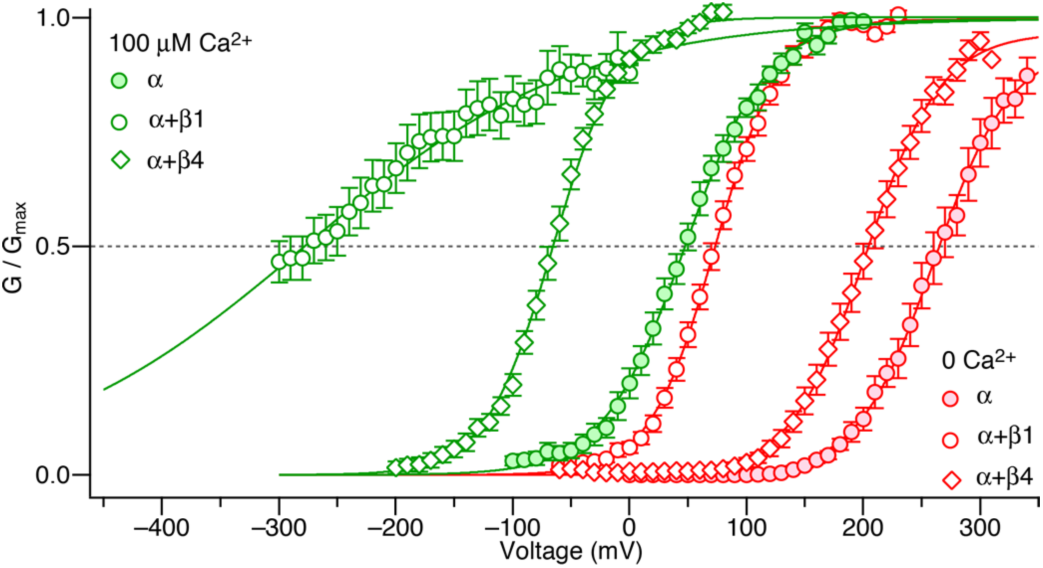
Ca^2+^-dependent activation of hSlo1 RKK:EEE channels. Normalized GV curves from hSlo1 RKK:EEE (filled circles), hSlo1 RKK:EEE+β1 (open circles) and hSlo1 RKK:EEE+β4 (open diamonds). The results without Ca^2+^ are shown in red and those with 100 µM Ca^2+^ are shown in green. n = 5 to 9. The smooth curves are Boltzmann-type fits (see Table S3).

**Figure S7.**
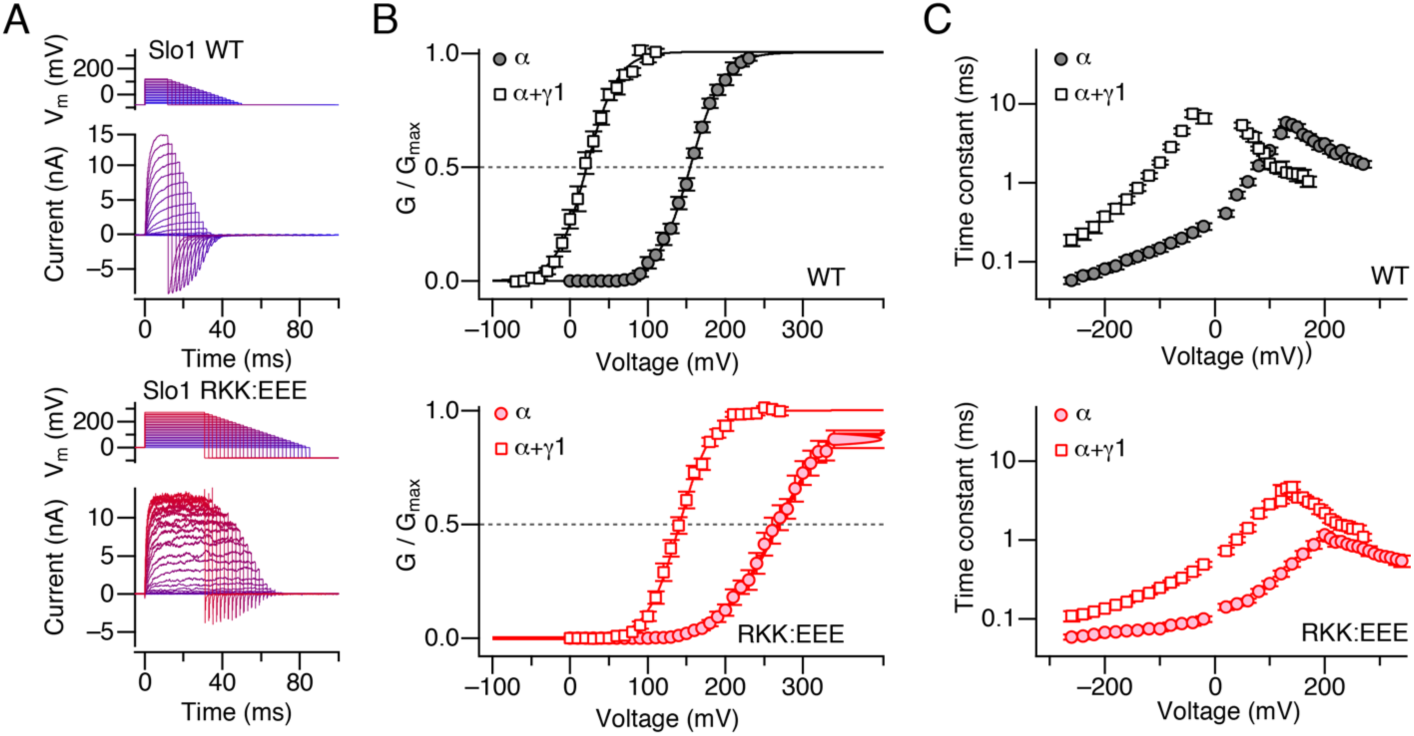
hSo1 RKK:EEE coassembles with γ1 (LRRC26). **A**. Illustrative currents from wild-type hSlo1+γ1 (*top*) and hSlo1 RKK:EEE+γ1 (*bottom*). **B**. Normalized GV curves from wild-type hSlo1 (*top* filled circles), wild-type hSlo1+γ1 (*top*, open squares), hSlo1 RKK:EEE (*bottom*, filled circles), and hSlo1 RKK:EEE+γ1 (*bottom*, squares). n = 6 to 9. The smooth curves are Boltzmann-type fits (Table S4). **C**. Time constants of current relaxation from wild-type hSlo1 (*top* filled circles), wild-type hSlo1+γ1 (*top*, squares), hSlo1 RKK:EEE (*bottom*, filled circles), and hSlo1 RKK:EEE+γ1 (*bottom*, squares). n = 6 to 9. All results were obtained without Ca^2+^.

**Figure S8.**
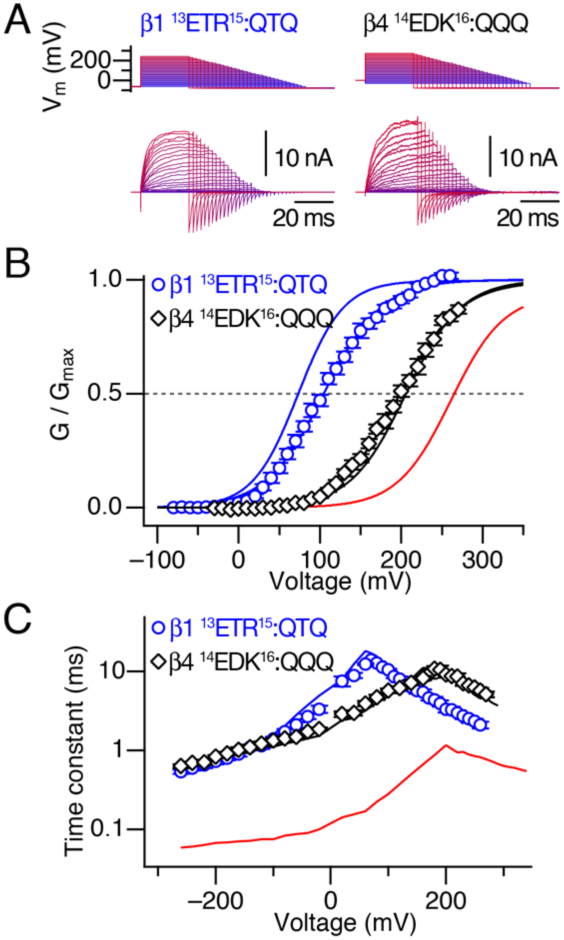
Electrophysiological characteristics of hSlo1 RKK:EEE coassembled with β1 E13Q:R15Q (^13^ETR^15^:QTQ) and β4 E14Q:D15Q:K16Q (^14^EDK^16^:QQQ). **A**. Illustrative currents from hSlo1 RKK:EE+β1 ^13^ETR^15^:QTQ (*left*) and β4 ^14^EDK^16^:QQQ (*right*). **B**. Normalized GV curves from the two channel types. The smooth curves are Boltzmann-type fits (see Table S5). **C**. Time constants of current relaxation from the two channel types. In each graph, the solid red curve represents the results from hSlo1 RKK:EEE alone, the solid blue curve represents those from hSlo1 RKK:EEE+β1, and the solid black curve represents those from hSlo1 RKK:EEE+β4.

**Figure S9.**
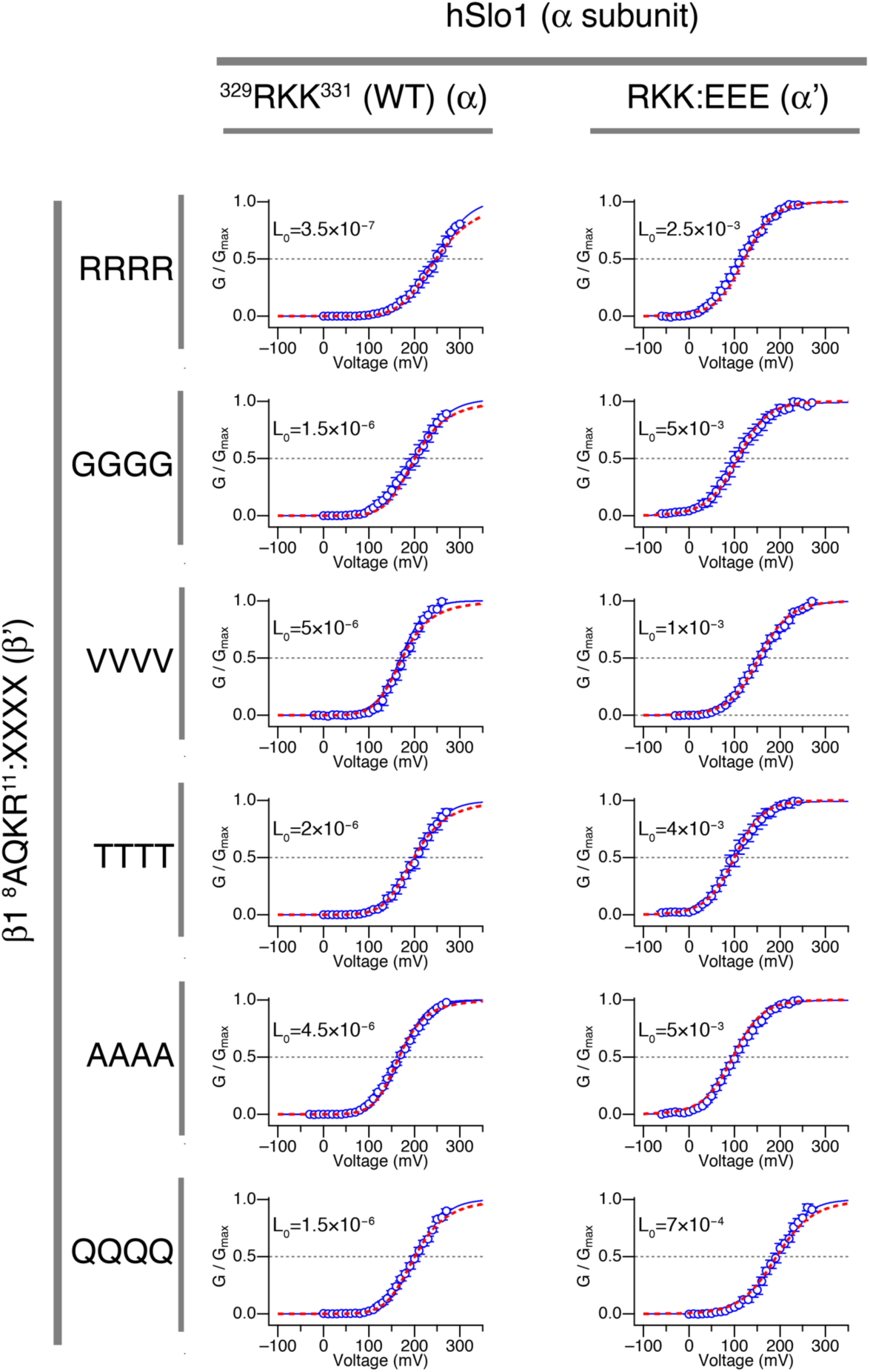
Electrophysiological properties of the mutants used in the double-mutant cycle analysis. The mutations analyzed are: hSlo1 WT (^329^RKK^331^) vs. hSlo1 RKK:EEE, and β1 wild-type (^8^AQKR^11^) vs. β1 AQKR:XXXX. In each GV graph, the red dashed curve represents the HA model prediction using the L_0_ value indicated.

**Figure S10.**
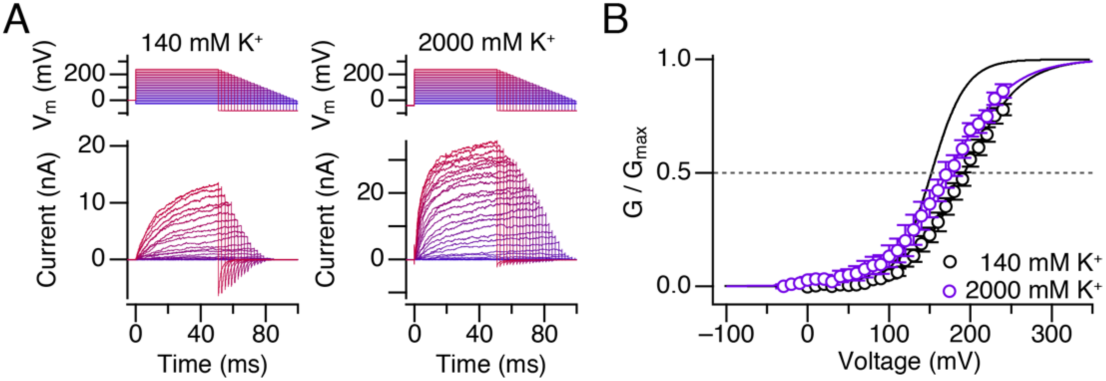
Increasing [KCl] to 2000 mM does not markedly alter hSlo1+β1 ^8^AQKR^11^: GGGG. **A**. Current traces with 140 mM or 2000 mM KCl inside from an illustrative patch. The voltage and current sweeps use the same color scheme, from hyperpolarized (blue) to depolarized (red) test voltages. **B**. Normalized GV curves with 140 and 2000 mM KCl. The black smooth curve the hSlo1 results. The smooth curves are Boltzmann-type fits (Table S7).

**Figure S11.**
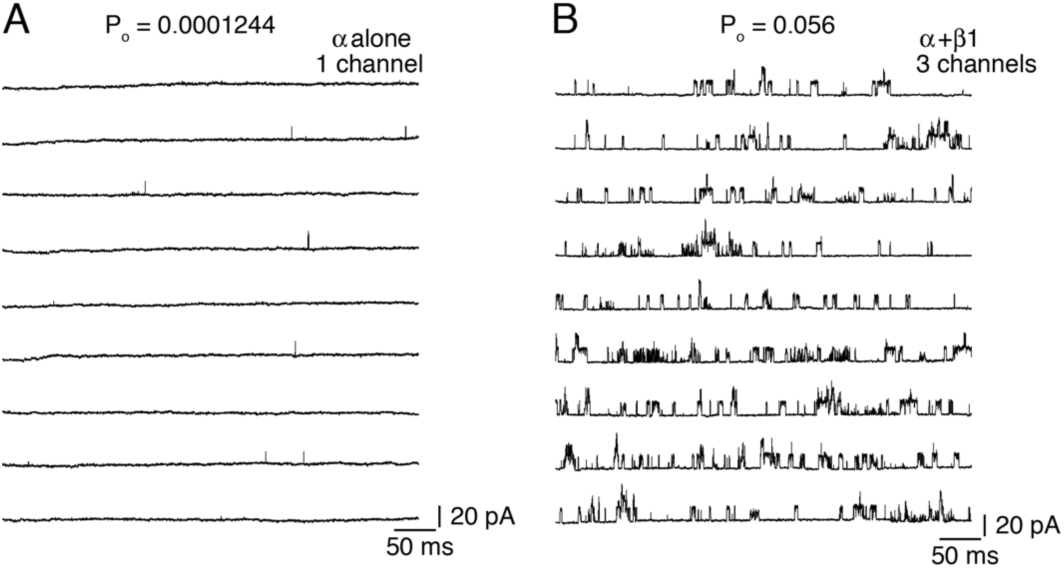
Single-channel openings from hSlo1 without the GR domain (hSlo1 ^329^RKK^331^:EEE ΔGR-Kv-minT) (31). **A**. Openings without β1. **B**. Openings with β1 coexpressed. The measurements were made at 120 mV and without Ca^2+^. Similar results were obtained in 2 patches.

**Figure S12.**
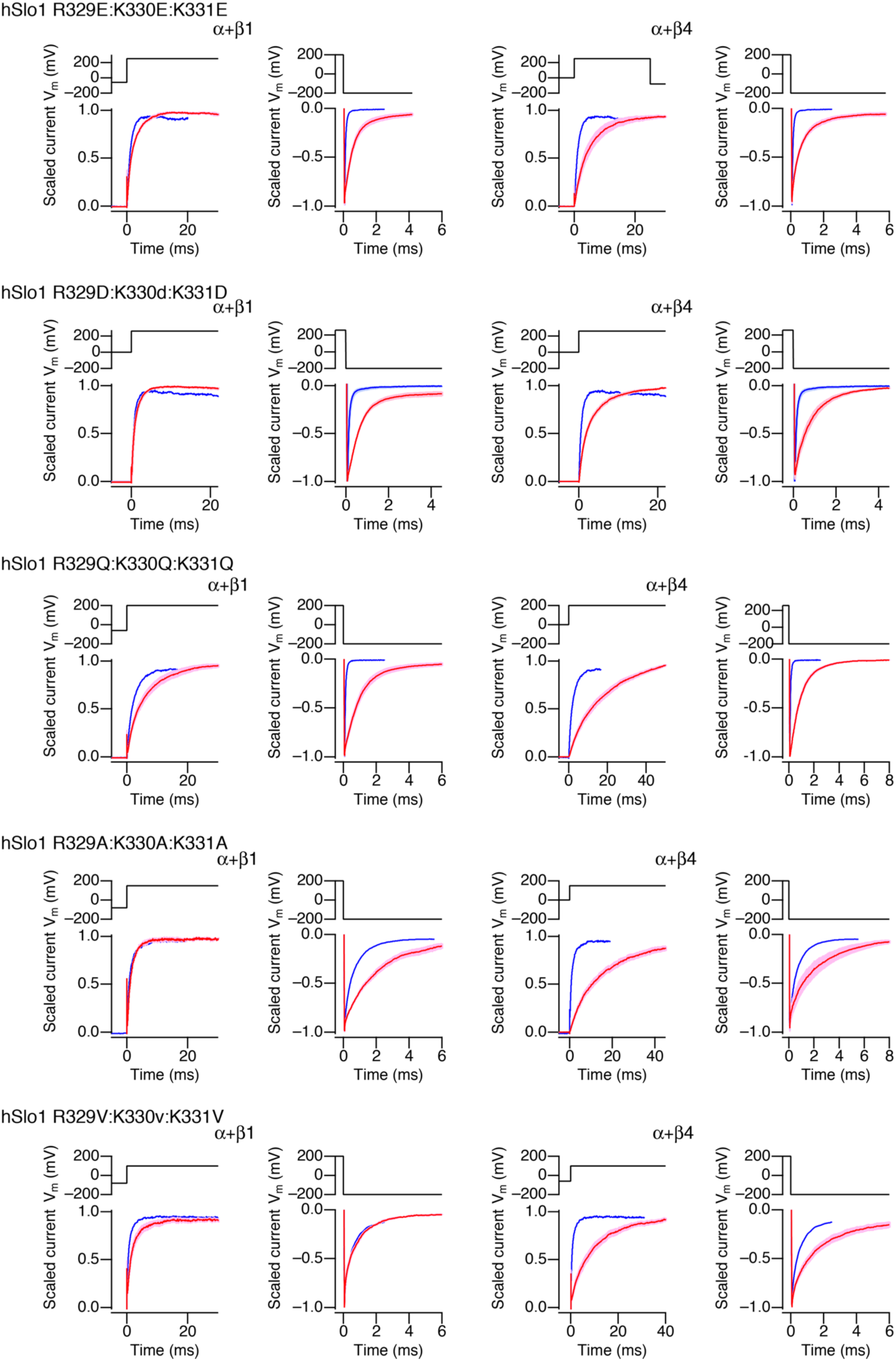
Changes in activation and deactivation time course by β1 and β4 in the hSlo1 RKK mutants. For each hSlo1 mutant, scaled currents at 200 mV and –200 mV from hSlo1 alone without any β (blue) and those with β1 or β4 (red) are shown. The trace width represents mean ± SEM. n = 7 to 15. All results were obtained without Ca^2+^.

**Figure S13.**
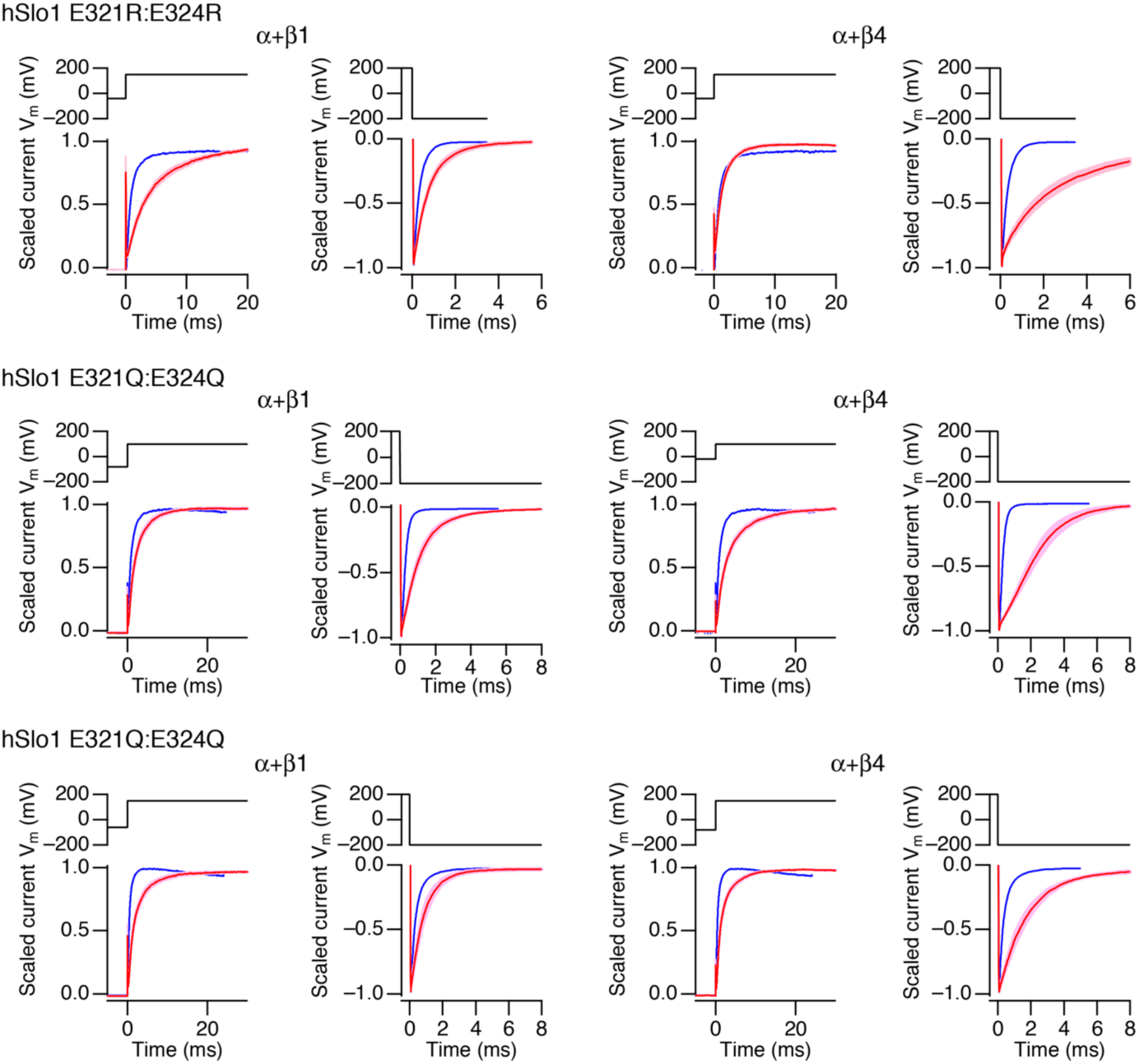
Changes in activation and deactivation time course by β1 and β4 in the hSlo1 E321/E324 mutants. Scaled currents at 200 mV and –200 mV from hSlo1 alone without any β (blue) and those with β1 or β4 (red) are shown. The sweep width represents mean ± SEM. n = 7 to 14. All results were obtained without Ca^2+^.

**Figure S14.**
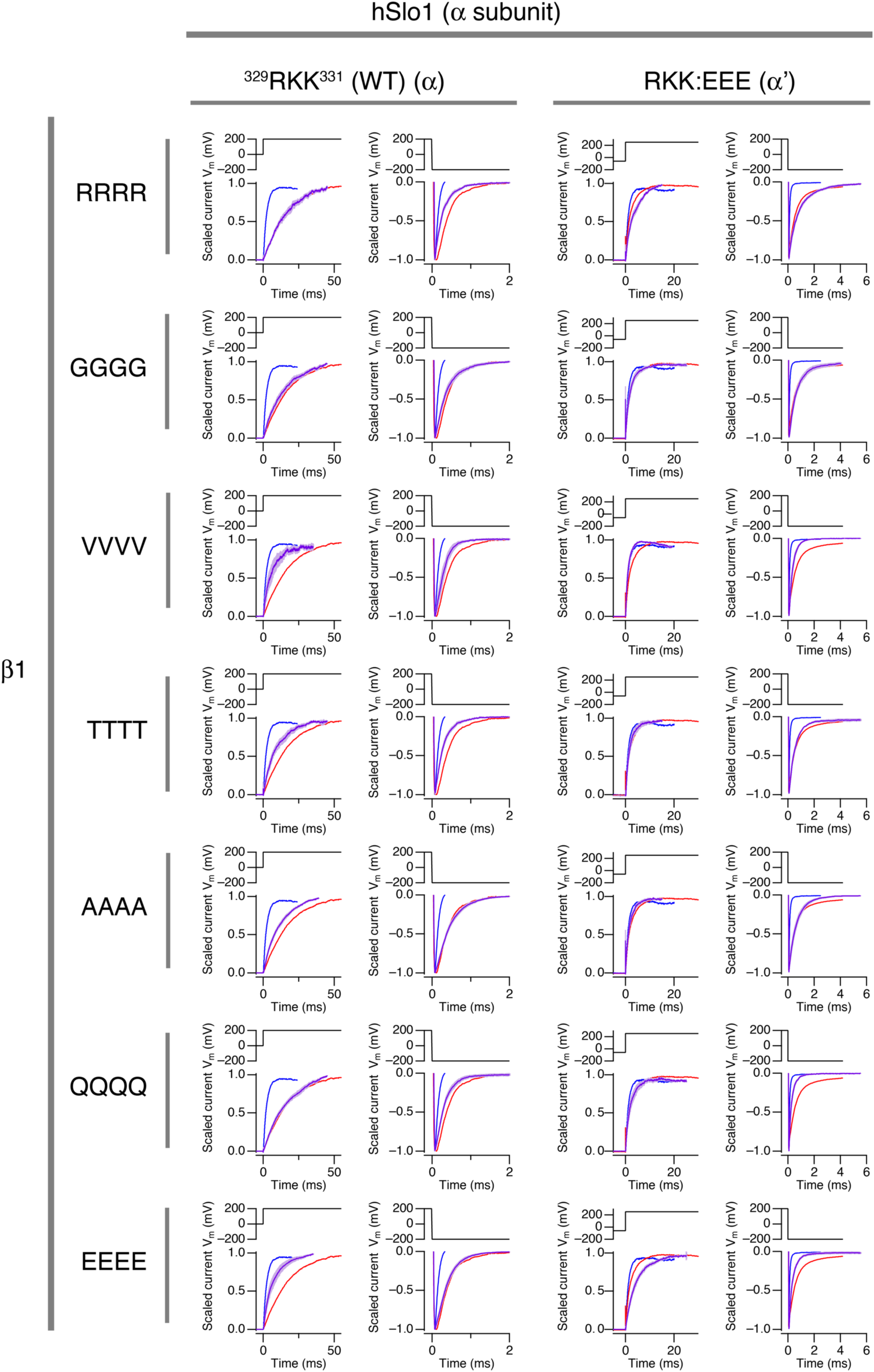
Changes in activation and deactivation time course by β1 and β4 in the Slo1 mutants used for the double-mutant analysis. Scaled currents at 200 mV and –200 mV. Blue: hSlo1 alone (*left*: WT, *right*: RKK:EEE). Red: hSlo1+ β1. Purple: hSlo1+mutant β1 indicated. The data trace width indicates mean ± SEM. n = 4 to 8. All results were obtained without Ca^2+^.

**Figure S15.**
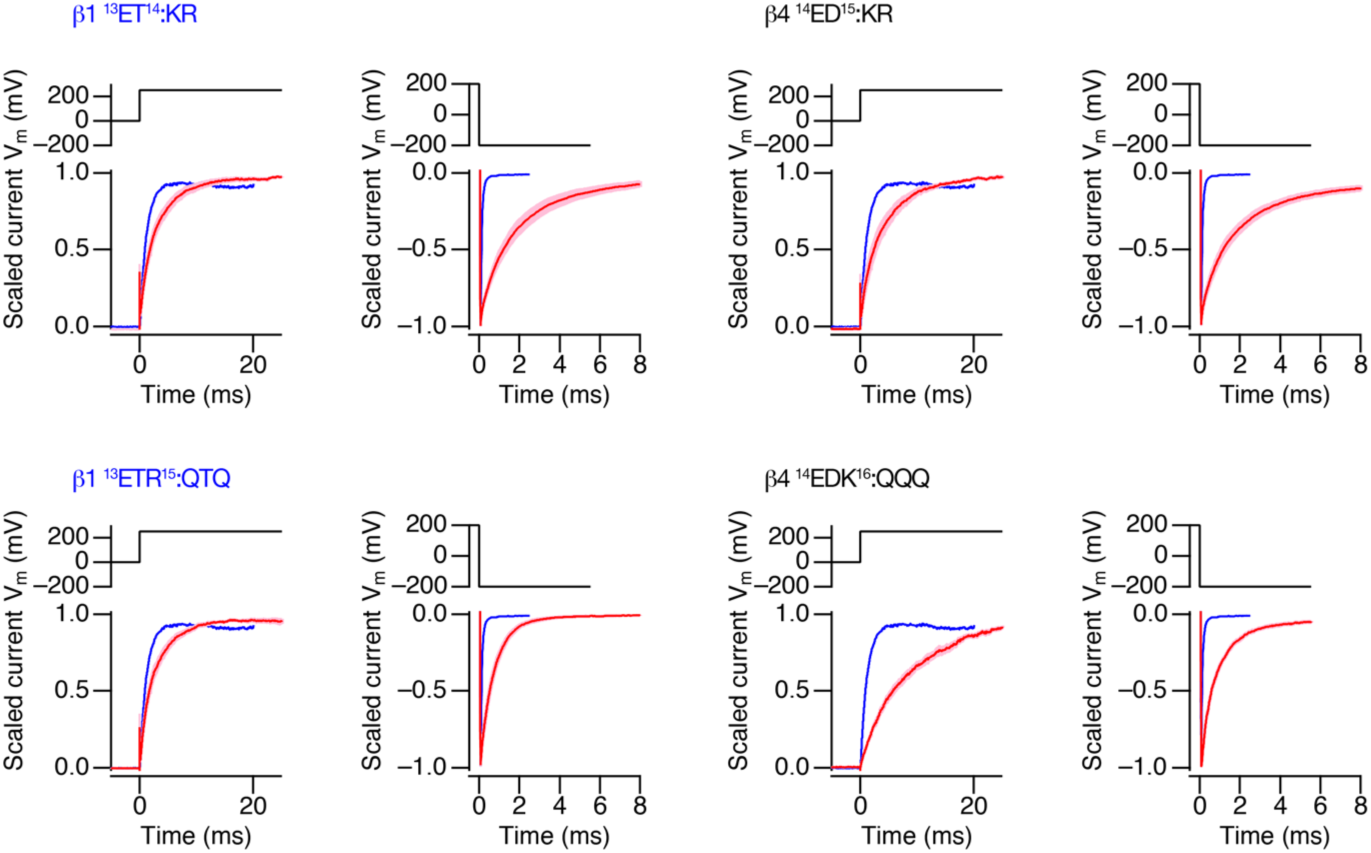
Changes in activation and deactivation time course by β1 E13Q:R15Q and β4 E14Q:D15Q:K16Q in hSlo1 RKK:QQQ. Blue: hSlo1 alone. Red: hSlo1+β1 E13Q:R15Q (*left*) and hSlo1+β4 E14Q:D15Q:R15Q (*right*). The data trace width indicates mean ± SEM. n = 6 to 9. All results were obtained without Ca^2+^.

**Figure S16.**
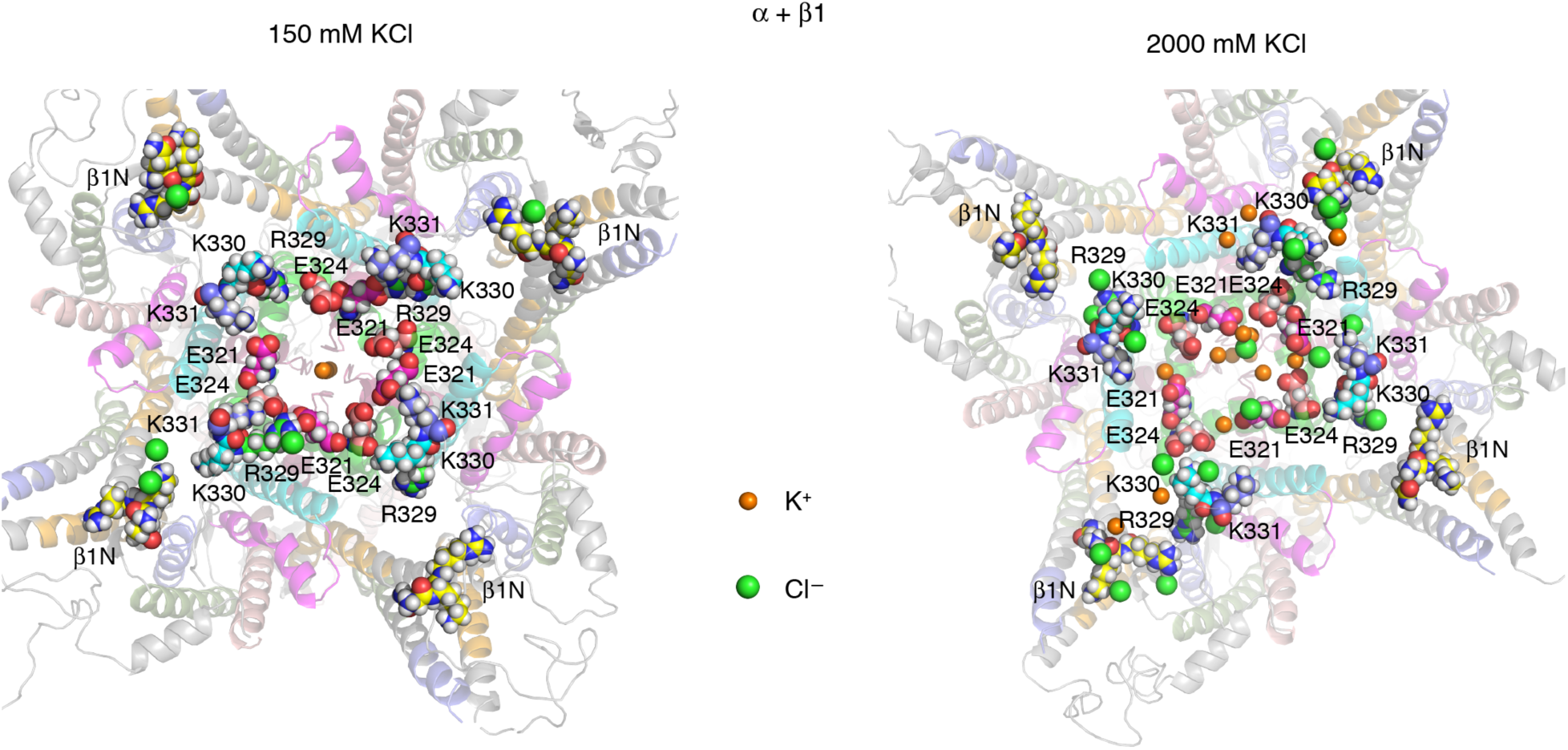
Comparison of wild-type hSlo1+β1 with 150 mM KCl (*left*) and 2000 mM KCl (*right*). Different residues are rendered using different carbon-color schemes; E321: magenta, E324: salmon, R329/E329: green, K330/E330: cyan, K331/E331: blue, β1 AQKR and β4 EYTE: yellow. The intracellular domain of the channel C-terminal to the RKK residues is not shown. Water, and lipids are not shown. Only those K^+^ and Cl^−^ ions within 5 Å of RKK and E321/E324 in addition to the pore K^+^ ions are shown.

**Table 1.** Activation characteristics of hSlo1 E321:E324 mutants coassembled with WT β1 and β4.

| Mutant | $V_{0.5}$ (mV) | $\Delta V_{0.5}$ (mV) | $Q_{app}$ | $Q_{app}$ ratio |
| --- | --- | --- | --- | --- |
| E321Q:E324Q | $85.7 \pm 9.7$ (13) | | $1.09 \pm 0.24$ (13) | |
| + $\beta 1$ | $21.6 \pm 13.8$ (10) | -64 | $0.95 \pm 0.14$ (13) | 0.87 |
| + $\beta 4$ | $80.7 \pm 10.0$ (10) | -5 | $1.16 \pm 0.16$ (13) | 1.06 |
| E321A:E324A | $79.4 \pm 8.2$ (16) | | $1.10 \pm 0.19$ (16) | |
| + $\beta 1$ | $48.6 \pm 10.2$ (12) | -30 | $1.05 \pm 0.14$ (12) | 0.95 |
| + $\beta 4$ | $85.8 \pm 20.3$ (14) | +5 | $1.02 \pm 0.18$ (14) | 0.93 |
| E321R:E324R | $92.9 \pm 11.5$ (15) | | $1.08 \pm 0.21$ (15) | |
| + $\beta 1$ | $74.8 \pm 19.7$ (12) | -18 | $0.74 \pm 0.18$ (12) | 0.69 |
| + $\beta 4$ | $39.1 \pm 25.2$ (18) | -54 | $0.83 \pm 0.23$ (18) | 0.77 |

Steady-state GV characteristics of the mutants with 0 $\text{Ca}^{2+}$ . The $\Delta V_{0.5}$ and $Q_{app}$ ratio values are in comparison with those of $\alpha$ alone. The results are mean ± standard deviation.

**Table 2.**
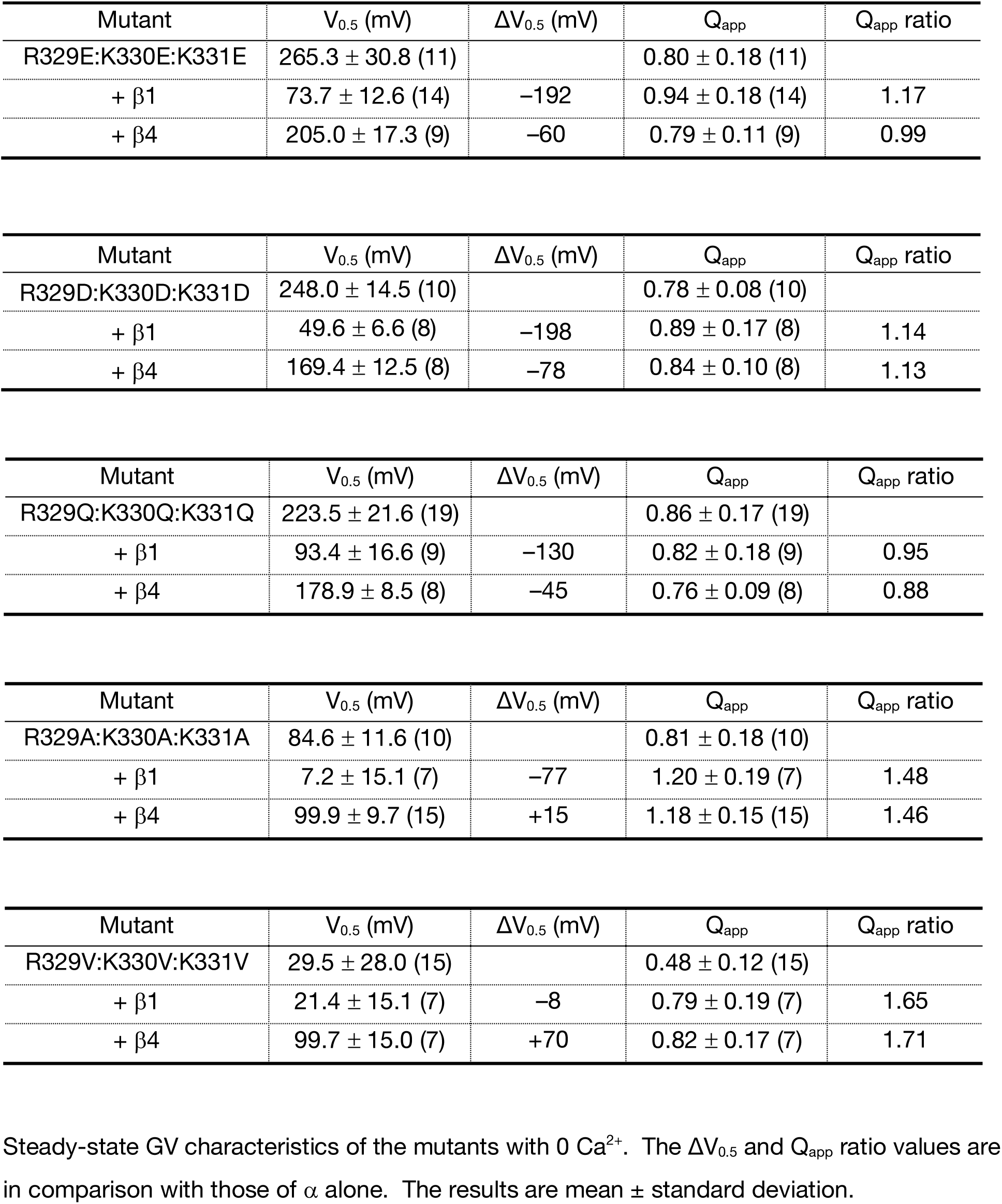
Activation characteristics of hSlo1 R329:K330:K331 mutants coassembled with β1 and β4.

**Table 3.** Activation characteristics of hSlo1 R329E:K330E:K331E coassembled with β1-β4 exchange mutants.

| Mutant | $V_{0.5}$ (mV) | $\Delta V_{0.5}$ (mV) | $Q_{app}$ | $Q_{app}$ ratio |
| --- | --- | --- | --- | --- |
| + β1 | $73.7 \pm 12.6$ (14) | | $0.94 \pm 0.18$ (14) | |
| + β1 R11E | $127.6 \pm 13.7$ (7) | +54 | $0.85 \pm 0.09$ (7) | 0.90 |
| + β1 K10T:R11E | $170.0 \pm 16.2$ (9) | +96 | $0.81 \pm 0.07$ (9) | 0.86 |
| + β1 A8E | $104.8 \pm 14.5$ (5) | +31 | $0.70 \pm 0.05$ (5) | 0.74 |
| + β1 AQKR:ETYE | $193.1 \pm 6.9$ (11) | +119 | $0.77 \pm 0.11$ (11) | 0.82 |
| + β4 | $205.0 \pm 17.3$ (9) | | $0.79 \pm 0.11$ (9) | |
| + β4 E12R | $128.4 \pm 4.7$ (9) | -77 | $1.08 \pm 0.15$ (9) | 1.36 |
| + β4 T11K:E12R | $103.5 \pm 17.9$ (12) | -101 | $1.03 \pm 0.29$ (12) | 1.30 |
| + β4 E9A | $166.9 \pm 28.3$ (11) | -38 | $0.84 \pm 0.09$ (11) | 1.06 |
| + β4 ETYE:AQKR | $83.1 \pm 10.0$ (11) | -122 | $0.80 \pm 0.13$ (11) | 1.01 |

Steady-state normalized GV characteristics of the mutants with 0 $\text{Ca}^{2+}$ . The $\Delta V_{0.5}$ and $Q_{app}$ ratio values are in comparison with those of WT $\beta$ 1 or β4. The results are mean ± standard deviation.

**Table 4.** Activation characteristics of hSlo1 R329E:K330E:K331E coassembled with β1 and β4 in the presence of 100 µM [Ca^2+^]_i_.

| Mutant | $V_{0.5}$ (mV) | $\Delta V_{0.5}$ (mV)* | $Q_{\text{app}}$ | $Q_{\text{app}}$ ratio |
| --- | --- | --- | --- | --- |
| R329E:K330E:K331E | $47.4 \pm 12.3$ (7) | -218 | $0.73 \pm 0.14$ (7) | 0.91 |
| + $\beta 1$ | $-257.8 \pm 27.0$ (5) | -332 | $0.28 \pm 0.12$ (5) | 0.30 |
| + $\beta 4$ | $-65.8 \pm 10.3$ (9) | -271 | $0.90 \pm 0.17$ (9) | 1.14 |

$\Delta V_{0.5}$ is calculated as $V_{0.5}(100 \mu\text{M} [\text{Ca}^{2+}]_i) - V_{0.5}(0 \text{ Ca}^{2+})$ .

**Table 5.**
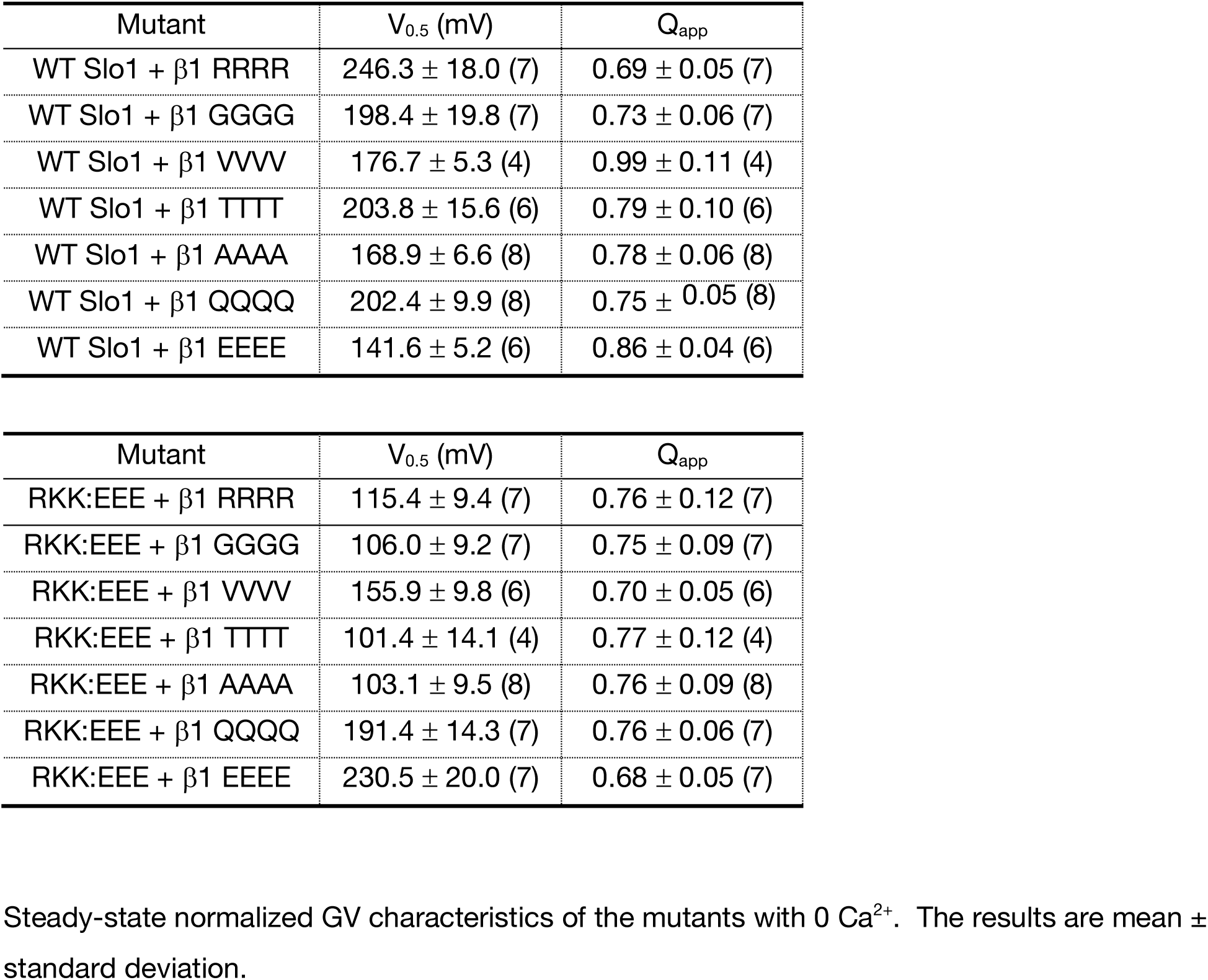
Activation characteristics of WT Slo1 and Slo1 R329E:K330E:K331E coassembled with β1 AQKR segment mutants.

**Table 6.**
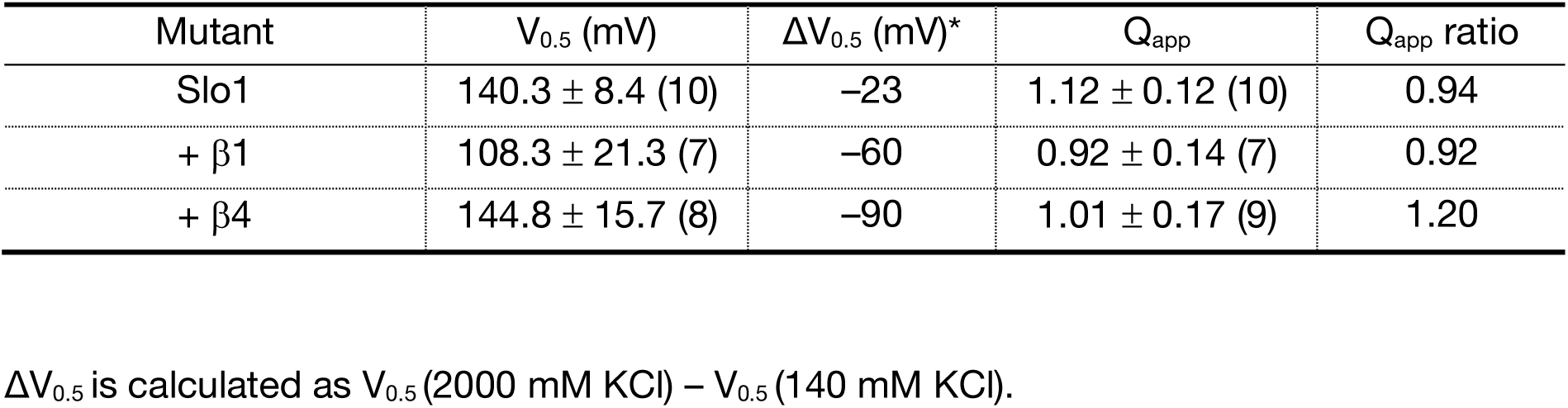
Activation characteristics of hSlo1 coassembled with β1 and β4 in the presence of 2000 mM KCl inside.

**Table 7.** Activation characteristics of hSlo1 R329E:K330E:K331E and WT hSlo1 coassembled with LRRC26 (γ1).

| Mutant | $V_{0.5}$ (mV) | $\Delta V_{0.5}$ (mV)* | $Q_{app}$ | $Q_{app}$ ratio |
| --- | --- | --- | --- | --- |
| R329E:K330E:K331E+ $\gamma 1$ | $140.9 \pm 7.2$ (6) | -125 | $1.24 \pm 0.13$ (6) | 1.55 |
| WT Slo1 + $\gamma 1$ | $20.8 \pm 10.4$ (9) | -140 | $1.38 \pm 0.18$ (9) | 1.2 |

$\Delta V_{0.5}$ is calculated as $V_{0.5}(\alpha + \gamma 1) - V_{0.5}(\alpha)$ .

